# The urinary proteome of individuals with a family history of frontotemporal dementia (FTD) but without known mutations differs from that of healthy controls

**DOI:** 10.1101/2024.11.25.625321

**Authors:** Yan Su, Min Chu, Haitong Wang, Liyong Wu, Youhe Gao

**Affiliations:** Gene Engineering Drug and Biotechnology Beijing Key Laboratory, College of Life Sciences, Beijing Normal University, Beijing 100875, China; Neurology, Xuanwu Hospital of Capital Medical University, Beijing 100053

**Keywords:** frontotemporal dementia, family history, urinary proteome

## Abstract

Frontotemporal dementia (FTD) is a common type of neurodegenerative dementia and the primary cause of dementia in individuals under 65 years old. Current studies typically consider family members carrying relevant gene mutations as genetic risk populations, while individuals with a family history of FTD but without known mutations are sometimes included as healthy controls in research, with their potential pathophysiological changes not being fully addressed. This study performed urinary proteomics analysis on individuals with a family history of FTD but without known mutations and healthy controls to investigate whether there are differential urinary proteins and biological pathways in FTD family members who remain undiagnosed with FTD or common mutations under existing diagnostic methods. The results showed 428 significantly differentially expressed proteins (P<0.01; FC≥2.0 or ≤0.5) between the two groups, among which multiple proteins have been reported to be involved in FTD or nervous system functions, particularly Progranulin (PGRN), recognized as an effective biomarker for FTD. The 146 enriched biological pathways (P<0.01) of differentially expressed proteins included multiple pathways directly related to neurons and the nervous system, such as glial cells. The differences in urinary proteins between individuals with a family history of FTD without known mutations and healthy controls indicate that urinary proteomics holds unique potential in exploring unknown pathogenic factors of FTD, providing a new perspective for deepening disease understanding and optimizing prevention and treatment strategies.

## 1 Introduction

### 1.1 Frontotemporal Dementia (FTD) and Its Research Status

Frontotemporal Dementia (FTD) is a progressive neurodegenerative disease that primarily affects the frontal and temporal lobes of the brain. These regions are responsible for controlling behavior, language, emotions, and decision-making abilities. Consequently, FTD patients exhibit impairments in cognitive and behavioral functions, language disorders, changes in behavior and emotions, impaired spatial orientation, and reduced judgment and decision-making abilities. Over time, these symptoms progress to more comprehensive dementia and may be complicated by other movement disorders such as amyotrophic lateral sclerosis^[1]^. FTD is a very common type of neurodegenerative dementia, predominantly affecting middle-aged and elderly individuals aged 45-65 years. It is the second most common form of dementia in individuals under 65 years of age, thus being recognized as an early-onset dementia^[2][3]^. The disease course ranges from 2 to 20 years, with an average of approximately 7-9 years, and the prognosis is poor^[4]^. Patients usually die from late-stage complications, including malnutrition, falls, and aspiration pneumonia^[5]^.

The etiology of FTD is not fully understood but may be related to genetic factors and gene mutations. Studies have shown that FTD has a strong genetic component, with approximately 30% of patients having a positive family history^[6]^. Mutations in the MAPT and progranulin (GRN) genes have been identified in approximately 50% of familial cases, while rare defects in the VCP, CHMP2B, TARDP, and FUS genes have been found in a small number of families^[7][8]^. Despite certain progress in research on the genetic mechanisms of FTD, there are currently few studies specifically targeting offspring who do not carry mutated genes. In many studies, these individuals are often used as control groups for affected or mutation-carrying patients within families.

Currently, there is no specific cure for FTD, and treatment mainly focuses on symptom management and exercise. Comprehensive treatment and nursing measures are employed to improve patients’ quality of life and delay disease progression, including pharmacological treatment (e.g., acetylcholinesterase inhibitors, anti-anxiety, and antidepressant drugs), rehabilitation therapy (e.g., cognitive training and functional training), behavioral therapy, nutritional therapy, family education, psychological counseling, and social participation. Early diagnosis and intervention can significantly improve the quality of life and prognosis of FTD patients ^[9][10][11]^_𰀬_

Clinical diagnosis of FTD is mainly based on neuropsychological assessments, neuroimaging examinations, and detection of cerebrospinal fluid (CSF) and plasma biomarkers^[12]^. Impaired executive function is a characteristic feature of FTD; however, in the early stages of the disease, executive dysfunction may be absent or masked by prominent behavioral changes. Additionally, since some types of FTD present symptoms similar to those in the early stage of Alzheimer’s disease (AD), such as episodic memory impairment, it is difficult to distinguish FTD from AD in the early stages using standard neuropsychological tests. Although existing neuroimaging methods can help distinguish FTD from AD, neuroimaging can detect abnormalities only when the disease has progressed to a certain extent with obvious structural and functional changes in the brain, thus failing to play a role in identification at the very earliest stage of the disease. Furthermore, due to pathological heterogeneity and significant variations in the severity of neurodegeneration, the ability of CSF biomarkers to reliably identify FTD is limited. Moreover, due to the regulation of the body’s homeostatic mechanisms, CSF and blood cannot detect the onset of the disease at an early stage^[13]^. Therefore, more sensitive detection methods are needed to achieve earlier diagnosis of FTD.

### 1.2 Urinary Biomarkers

The urinary proteome holds great potential as a next-generation biomarker. Firstly, urine is not regulated by the body’s homeostatic mechanisms, enabling it to sensitively reflect early physiological or pathological changes in the body, thus playing an important role in the early diagnosis of diseases^[14]^. Secondly, urine is easy to collect, preserve, and analyze; long-term cryopreservation can be achieved by directly binding urinary proteins to PVDF membranes^[15]^. These characteristics make urine a highly promising biomarker, allowing it to maximize its role in physiological and pathophysiological research.

In studies on urinary proteomics, the use of animal models facilitates variable control and minimizes interference from relevant factors, enabling sample collection at the very earliest stages of disease onset and progression^[16][17]^. Multiple studies using animal models have shown that urinary proteomes can be detected earlier than changes in blood, symptoms, signs, imaging, and pathology for disease diagnosis. Currently, research on urinary proteomes in various diseases affecting multiple organs (such as the liver, kidney, brain, and lung), including tumors, degenerative diseases, and inflammation, has already verified the potential of urinary proteomes in early diagnosis^[18][13][19][20][21][22][23]^.

Urinary proteomics has been applied to the study of various human diseases and has demonstrated great potential in the search for early diagnostic biomarkers for certain diseases, with broad prospects in future clinical applications. Rubio-Sánchez R et al. ^[24]^ were the first to investigate microorganisms present in vaginal secretions and urine of patients infected with Trichomonas vaginalis to identify volatile biomarkers. They found that volatile compoundgs such as 2-octen-1-ol and 3-nonanone could serve as biomarkers in urine samples, and 3-hydroxy-2,4,4-trimethylpentyl 2-methylpropanoate, identified in vaginal secretions, was highly correlated with urine from patients with trichomoniasis. Additionally, exploring changes in the urinary proteome is an important approach for early detection of diagnostic markers for urinary system diseases and some systemic malignancies, particularly shining in studies on bladder cancer. Previous studies have identified numerous potential markers for transitional cell carcinoma, including hemoglobin, tumor-associated trypsin inhibitor, soluble E-cadherin, and hyaluronidase, some of which have been used in clinical practice^[25]^.

Therefore, this study conducts urinary proteomic analysis on individuals with a family history of FTD who do not carry known mutations and healthy individuals, aiming to investigate whether individuals who have not been identified with FTD or common mutations using existing diagnostic methods have urinary biomarkers and biological pathways that differ from those of healthy individuals. This research seeks to deepen the understanding of the disease and provide a basis and clues for the prevention and treatment of FTD.

## 2 Materials and Methods

### 2.1 Urine Sample Collection

This study was approved by the Ethics Committee of Xuanwu Hospital (approval No. Linyan shen ^[2023]^ 036). All participants provided written informed consent. Participants were recruited from Xuanwu Hospital, and samples were collected on August 26, 2023. The participant samples included 5 samples from individuals with a family history of FTD with MAPT gene mutation but without carrying MAPT or other gene mutations, 1 sample from a MAPT gene mutation carrier, and 1 sample from a FTD patient with MAPT gene mutation. The collected urine samples were stored in a -80°C refrigerator. Healthy control samples were obtained from a published preprint study on COVID-19, with consistent sample processing methods as in this study, totaling 11 cases^[26]^. This study mainly compared the urinary proteomes between individuals with a family history of FTD but without known mutations and healthy controls; the MAPT gene mutation carrier and the FTD patient with MAPT gene mutation were only used for cluster analysis.

### 2.2 Urine Sample Processing

Urinary proteins extraction: After thawing the urine samples, centrifuge at 12000 × g and 4°C for 20 minutes, and transfer the supernatant to a new EP tube. Then add dithiothreitol solution (DTT, Sigma) to a final concentration of 20 mM, heat in a metal bath at 37°C for 60 minutes, and cool to room temperature. Then add iodoacetamide (IAA, Sigma) to a final concentration of 50 mM, and place in the dark at room temperature for 40 minutes. Add four times the volume of absolute ethanol, mix well, and precipitate in a - 20°C refrigerator for 24 hours. The next day, centrifuge the mixture at 12000 × g and 4 °C for 30 minutes, discard the supernatant, and resuspend the protein precipitate in lysis buffer (containing 8 mol/L urea, 2 mol/L thiourea, 25 mmol/L DTT, 50 mmol/L). Centrifuge again at 12000 × g and 4°C for 30 minutes, and retain the supernatant. Measure the protein concentration using the Bradford method.

Protease digestion: Take 100 μg of the urinary proteins sample into an EP tube, add 25 mmol/L NH4HCO3 solution to make the total volume 200 μL as a sample for standby. Next, wash the membrane of the 10 kDa ultrafiltration tube (Pall, Port Washington, NY, USA): first add 200 μL of UA solution (8 mol/L urea, 0.1 mol/L Tris - HCl, pH 8.5), centrifuge at 14000 × g and 18°C for 10 minutes, and repeat twice. Add the above - prepared sample to the ultrafiltration tube and centrifuge at 14000 × g and 18°C for 40 minutes. Then add 200 μL of UA solution and centrifuge at 14000 × g and 18°C for 30 minutes, and repeat twice. Subsequently, add 25 mmol/L NH4HCO3 solution and centrifuge at 14000 × g and 18°C for 30 minutes, and repeat twice. Finally, add trypsin (Trypsin Gold, Promega, Fitchburg, WI, USA) at a ratio of trypsin:protein = 1:50 and incubate overnight at 37°C. After overnight incubation, centrifuge to collect the digested filtrate, perform desalting treatment through an Oasis HLB solid - phase extraction column, and vacuum - dry to obtain peptide lyophilization, which is stored at - 80°C.

### 2.3 LC - MS/MS Tandem Mass Spectrometry Analysis

Reconstitute the peptide with 0.1% formic acid water and dilute the peptide concentration to 0.5 μg/μL. Separate the peptides through a Thermo ESAY - Nlc1200 liquid chromatography system with the parameter settings as follows: elution time of 90 minutes and elution gradient (A phase: 0.1% formic acid; B phase: 80% acetonitrile). Repeat data - dependent mass spectrometry data acquisition on the separated peptides three times through an Orbitrap Fusion Lumos Tribird mass spectrometer using the data - independent acquisition mode.

### 2.4 Database Search

Use the Spectronaut Pulsar (Biognosys AG, Switzerland) software to search the database of the raw files acquired by the mass spectrometer and compare them with the SwissProt Human database. Calculate the peptide abundance by adding the peak areas of the respective fragment ions in MS2. Calculate the protein abundance by adding the abundances of the respective peptides.

### 2.5 Statistical and Bioinformatics Analysis

Each sample was technically repeated three times, and the average value of the obtained data was used for statistical analysis. This experiment performed a group comparison between the FTD family history unaffected group and the healthy group to screen for differential proteins. The screening conditions for differential proteins were as follows: the fold change (FC) between groups ≥ 2 or ≤ 0.5, and the P value of the two - tailed unpaired t - test analysis < 0.01. The screened differential proteins were queried for their names and functions through the Uniprot website (https://www.uniprot.org), and the biological function enrichment analysis was performed through the DAVID database (https://david.ncifcrf.gov). The reported literature was retrieved in the Pubmed database (https://pubmed.ncbi.nlm.nih.gov) to analyze the functions of the differential proteins.

## 3 Results

### 3.1 Random Grouping

Samples from the healthy group (n=11) and the group with FTD family history without known mutations (n=5) were randomly divided into two groups, resulting in a total of 4368 possible groupings. Among all random combination types, the average number of differential proteins across all randomizations was calculated using the same screening criteria. The ratio of this average number to the number of differential proteins obtained from the normal grouping was defined as the proportion of randomly generated differential proteins. The results of random grouping showed that the proportion of randomly generated differential proteins was 4.24%, which was less than 5%. This indicates that most of the differential proteins identified in this study were not randomly generated, confirming that there are significant differences in the urinary proteomes between the group with FTD family history without known mutations and the healthy group.

### 3.2 Comparison of Urinary Proteomes Between the Group with FTD Family History Without Known Mutations and the Healthy Group

#### 3.2.1 Differential Proteins

Urinary proteins from the group with FTD family history without known mutations and the healthy group were compared. The criteria for screening differential proteins were: fold change (FC) value (the ratio of the number of proteins identified in urine samples from individuals with FTD family history without known mutations to that in urine samples from healthy individuals) ≥ 2 or ≤ 0.5, and P < 0.01 by two-tailed unpaired t-test. The results showed that, compared with the healthy group, a total of 428 differential proteins were identified in the group with FTD family history without known mutations, including 399 downregulated proteins and 29 upregulated proteins. These differential proteins were sorted by FC in ascending order, and their detailed information, retrieved via Uniprot, is listed in Supplementary Table 1. Information on the top 20 most significantly upregulated and downregulated differential proteins is presented in Table 2.

**Table 1.** Information and statistical results of collected samples.

| Sample information |  | Healthy (n=11) | Individuals with FTD family history without MAPT or other gene mutations (n=5) | FTD patient with MAPT gene mutation (n=1) | MAPT gene mutation carrier (n=1) |
| --- | --- | --- | --- | --- | --- |
| Age | Mean value | 42.9±6.5 | 45.4±10.4 | 33±0 | 59±0 |
|  | Age range | 34-56 | 33-61 | 33 | 59 |

**Table 2.** Information on the top 20 most significantly upregulated and downregulated differential proteins screened with FC ≥ 2.0 or ≤ 0.5 and P < 0.01 between the group with FTD family history without known mutations and the healthy group.

| Uniprot ID | Protein names | Fold change | P value | Trend |
| --- | --- | --- | --- | --- |
| Q13467 | Frizzled-5 | 0.00 | 2.57E-05 | ↓ |
| P05106 | Integrin beta-3 | 0.00 | 2.08E-04 | ↓ |
| Q6UXH9 | Inactive serine protease PAMR1 | 0.00 | 4.82E-04 | ↓ |
| Q9NRR1 | Cytokine-like protein 1 | 0.00 | 1.17E-03 | ↓ |
| Q9NQ34 | Transmembrane protein 9B | 0.00 | 1.47E-03 | ↓ |
| Q13705 | Activin receptor type-2B | 0.00 | 2.08E-03 | ↓ |
| Q7Z4P5 | Growth/differentiation factor 7 | 0.00 | 3.11E-03 | ↓ |
| P30486 | HLA class I histocompatibility antigen, B alpha chain | 0.00 | 5.57E-03 | ↓ |
| P26012 | Integrin beta-8 | 0.00 | 3.22E-05 | ↓ |
| A0A0C4DH35 | Probable non-functional immunoglobulin heavy variable 3-35 | 0.00 | 3.90E-06 | ↓ |
| P01782 | Immunoglobulin heavy variable 3-9 | 0.00 | 9.33E-05 | ↓ |
| A0A087WW87 | Immunoglobulin kappa variable 2-40 | 0.00 | 1.75E-03 | ↓ |
| Q86UP0 | Cadherin-24 | 0.00 | 3.19E-03 | ↓ |
| P23434 | Glycine cleavage system H protein, mitochondrial | 0.01 | 3.48E-04 | ↓ |
| P0DP01 | Immunoglobulin heavy variable 1-8 | 0.01 | 3.84E-05 | ↓ |
| O75581 | Low-density lipoprotein receptor-related protein 6 | 0.02 | 4.04E-03 | ↓ |
| Q01628 | Interferon-induced transmembrane protein 3 | 0.02 | 9.17E-05 | ↓ |
| Q7KYR7 | Butyrophilin subfamily 2 member A1 | 0.02 | 4.14E-04 | ↓ |
| P35556 | Fibrillin-2 [Cleaved into: Placensin] | 0.02 | 6.30E-03 | ↓ |
| O75015 | Low affinity immunoglobulin gamma Fc region receptor III-B | 0.03 | 1.44E-03 | ↓ |
| P00492 | Hypoxanthine-guanine phosphoribosyltransferase | 2.64 | 8.18E-03 | ↑ |
| Q9UMY4 | Sorting nexin-12 | 2.83 | 3.69E-03 | ↑ |
| Q9Y3B8 | Oligoribonuclease, mitochondrial | 2.85 | 2.92E-03 | ↑ |
| P84095 | Rho-related GTP-binding protein RhoG | 2.88 | 5.48E-03 | ↑ |
| Q9H3Z4 | DnaJ homolog subfamily C member 5 | 2.97 | 2.43E-03 | ↑ |
| O43681 | ATPase GET3 | 3.43 | 3.45E-03 | ↑ |
| P12081 | Histidine--tRNA ligase, cytoplasmic | 3.45 | 1.44E-04 | ↑ |
| Q9H3K6 | BolA-like protein 2 | 3.45 | 7.38E-03 | ↑ |
| P14138 | Endothelin-3 | 3.52 | 7.04E-03 | ↑ |
| P52943 | Cysteine-rich protein 2 | 3.79 | 1.44E-03 | ↑ |
| Q8NFP4 | MAM domain-containing glycosylphosphatidylinositol anchor protein 1 | 4.13 | 5.30E-04 | ↑ |
| Q13261 | Interleukin-15 receptor subunit alpha | 4.40 | 2.93E-03 | ↑ |
| P54577 | Tyrosine--tRNA ligase, cytoplasmic | 4.73 | 2.11E-03 | ↑ |
| Q8N5J2 | Ubiquitin carboxyl-terminal hydrolase MINDY-1 | 5.95 | 1.74E-06 | ↑ |
| P12532 | Creatine kinase U-type, mitochondrial | 6.77 | 9.66E-05 | ↑ |
| O00534 | von Willebrand factor A domain-containing protein 5A | 7.25 | 1.96E-04 | ↑ |
| Q2TAA2 | Isoamyl acetate-hydrolyzing esterase 1 homolog | 10.28 | 8.46E-03 | ↑ |
| P35247 | Pulmonary surfactant-associated protein D | 15.09 | 1.51E-03 | ↑ |
| P16949 | Stathmin | 18.26 | 2.32E-03 | ↑ |
| P62857 | Small ribosomal subunit protein eS28 | 21.29 | 2.95E-03 | ↑ |

#### 3.2.2 Functional Analysis of Differential Proteins

The identified differential proteins were searched in the PubMed database.

Among them, 8 differential proteins showed the most significant downregulation, with changes “from presence to absence”. These proteins are Frizzled-5, Integrin beta-3, Inactive serine protease PAMR1, Cytokine-like protein 1, Transmembrane protein 9B, Activin receptor type-2B, Growth/differentiation factor 7, and HLA class I histocompatibility antigen (B alpha chain). Proteins that have been reported to be related to neurological diseases or physiological processes of the nervous system are as follows:

① Integrin beta-3: Neuroinflammation involves cytokine release, astrocyte reactivity and migration. Neuronal Thy-1 can promote astrocyte migration by binding to Integrin beta-3 and syndecan-4^[27]^.

② Frizzled-5: Frizzled-5 is a receptor for Wnt proteins (WNT2, WNT10B, WNT5A). Studies have found that Frizzled 5 is continuously and cell-autonomously required for the survival of adult paraventricular nucleus neurons, and defects in the Wnt-Frizzled signaling pathway may be the cause of neuronal loss in degenerative central nervous system diseases^[28]^.

③ Activin receptor type-2B: During the development of chicken ciliary ganglia, researchers found that signaling through activin receptor 2B plays an important role in regulating cell death during nervous system development. Downregulation of its expression can lead to dysregulation of neuronal apoptosis, affect programmed cell death in neurodevelopment, and hinder the differentiation of choroidal neurons^[29]^.

④ Growth/differentiation factor 7 (GDF7): GDF7 is expressed in the rhombic lip of the cerebellum and plays a key role in cerebellar development. GDF7 lineage cells contribute to various neuronal and glial cell types in the cerebellum^[30]^.

In addition, among other downregulated proteins, there are also proteins related to the nervous system. Cadherin-24 (FC=0.0044, P=3.19E-03): It has been found that presenilin 1 can form a complex with N-cadherin, maintaining intercellular junctions and neurite outgrowth^[31]^. Glycine cleavage system H protein, mitochondrial (FC=0.0055, P=3.48E-04): This protein plays a key role in glycine metabolism, and glycine is an important neurotransmitter involved in inhibitory neurotransmission^[32]^. Low-density lipoprotein receptor-related protein 6 (FC=0.017, P=4.04E-03): The expression level of LRP6 is significantly decreased in the brains of Alzheimer’s disease (AD) patients, and loss of LRP6 function can increase inflammatory markers, mitochondrial dysfunction, and stroke volume^[33]^. Interferon-induced transmembrane protein 3 (FC=0.018, P=9.17E-05): Its upregulation can lead to increased production of β-amyloid protein. In addition, the expression of this protein in cerebrovascular endothelial cells is also related to the pathogenesis of Alzheimer’s disease, and inhibiting the expression of interferon-induced transmembrane protein 3 can improve AD pathology and cognitive impairment^[34]^. Low affinity immunoglobulin gamma Fc region receptor III-B (Fc-γRIII-β) (FC=0.026, P=1.44E-03): It is one of the low-affinity immunoglobulin gamma Fc region receptors. Fcγ receptors (including Fcγ RIII-B) are expressed in microglia and astrocytes and participate in immune responses in the central nervous system; abnormal activation of these receptors may lead to neuroinflammation, thereby affecting the progression of neurodegenerative diseases^[35]^.

Among the upregulated differential proteins, Stathmin (FC=18.26, P=2.32E-03) has increased expression during neuronal differentiation and plasticity and is altered in many neurodegenerative diseases^[36]^. Creatine kinase U-type, mitochondrial (FC=6.77, P=9.66E-05): Bioenergetic dysfunction and mitochondrial damage are directly or indirectly related to the pathogenesis of many neurodegenerative diseases. The creatine kinase/phosphocreatine system, where creatine kinase is located, plays an indispensable role in energy buffering and overall cellular bioenergetics; therefore, abnormalities in this system may exacerbate the development of neurodegenerative diseases^[37]^. Interleukin-15 receptor subunit alpha (FC=4.40, P=2.93E-03) can regulate the self-renewal of neural stem cells and neurogenesis. Its role in regulating neuroinflammation and microglial activation suggests that it may be related to the pathological mechanism of AD^[38]^. MAM domain-containing glycosylphosphatidylinositol anchor protein 1 (FC=4.40, P=2.93E-03) and its MAM domain play an important role in cognitive function in mice, especially memory related to novel object recognition. This role may be mediated by the regulation of synaptic inhibition in the hippocampus by MDGA1, thereby affecting the function of neural circuits and ultimately cognitive behavior^[39]^. Endothelin-3 (FC=3.52, P=7.04E-03): Studies have shown that endothelin increases the number of reactive astrocytes involved in Alzheimer’s disease^[40]^. Rho-related GTP-binding protein RhoG (FC=2.88, P=5.48E-03) is a member of the Rho GTPases family. Abnormal expression or dysfunction of Rho GTPases may lead to neurodevelopmental disorders, such as intellectual disability and motor dysfunction^[41]^. Sorting nexin-12 (FC=2.88, P=5.48E-03) can indirectly affect the β-processing of amyloid precursor protein to produce β-amyloid through their interaction, and reduced levels of this protein have been found in the brains of Alzheimer’s disease patients^[42]^. Deficiency of Hypoxanthine-guanine phosphoribosyltransferase (FC=2.88, P=5.48E-03) can affect an individual’s neurobehavior, which is related to dysfunction of the dopaminergic pathway in the basal ganglia, thereby affecting neurodevelopment-related processes such as neurite outgrowth^[43]^.

Notably, Progranulin (PGRN) (FC=0.27, P=7.70E-04) was identified among the differential proteins. PGRN is encoded by the GRN gene, is a secreted protein, and exerts its functions through intercellular signal transmission and signal transduction, being related to cell proliferation, wound repair, and growth factors. Multiple studies have shown that GRN mutations can lead to PGRN haploinsufficiency, ultimately causing FTD, and PGRN is considered an effective biomarker^[8][44][45]^. GRN mutations may affect its normal function by reducing PGRN expression, secretion, and altering subcellular localization^[46]^. Studies have found that decreased plasma PGRN levels can predict GRN mutations^[47]^.

In addition, there are other reported proteins related to FTD or neurological diseases. Vascular endothelial growth factor A, long form (L-VEGF) (FC=0.27, P=5.12E-03): The expression level of VEGF is significantly decreased in the brains of AD patients, which may play an important role in the pathogenesis of AD, and changes in its level are related to disease progression^[48]^. We also found changes in Metalloproteinase inhibitor 3 (TIMP-3) (FC=0.063, P=4.32E-03). Although there are no reports on the relationship between TIMP-3 and neurological diseases, studies have shown that the levels of its family members TIMP-1 and TIMP-2 are altered in FTD patients^[49][50]^.

Our study shows that among the differential proteins between the group with FTD family history without known mutations and the healthy group, a large number of significantly changed proteins have been reported to be related to FTD or the nervous system. In addition, some significantly changed differential proteins have not been reported to be related to FTD or the nervous system. Further research on the functions of these proteins in FTD or the nervous system may help identify early screening biomarkers or potential drug targets for FTD.

#### 3.2.3 Enrichment Analysis of Biological Processes for Differential Proteins

The DAVID database was used to perform enrichment analysis of biological processes (BP) for the 428 differential proteins identified by group analysis. The results showed that a total of 146 BP pathways were significantly enriched (P < 0.01), with detailed information listed in Supplementary Table 2.

Among the 5 biological pathways with the lowest enrichment P-values, 3 were related to adhesion, namely “cell adhesion”, “heterophilic cell-cell adhesion via plasma membrane cell adhesion molecules”, and “homophilic cell adhesion via plasma membrane adhesion molecules”. Cell adhesion molecules (CAMs) are involved in the formation and maintenance of synapses between neurons, as well as the regulation of synaptic plasticity. In patients with Alzheimer’s disease, changes in the levels and functions of CAMs can affect synaptic plasticity, thereby impairing signal transmission between neurons and the function of neural networks; changes in CAMs may also affect the migration and activation of inflammatory cells, further influencing the progression of neuroinflammation^[51]^. The other 2 pathways among the 5 with the lowest P-values were “angiogenesis” and “blood coagulation”. Cerebral microbleeds are common in neurocognitive disorders and have been found to increase the risk of dementia, including Alzheimer’s disease, which explains the presence of these two biological processes^[52]^.

Meanwhile, we enriched many pathways directly related to neurons and the nervous system, as shown in Figure 1a. Multiple studies have indicated that glial cells are key pathogenic factors in FTD^[53][54]^. Glial cells include various types such as astrocytes and oligodendrocytes, which provide support and protection for neurons, and participate in the development, repair, and immune response of the nervous system. Astrocytes may be involved in the progression of FTD by releasing inflammatory factors, and may also participate in the process of abnormal protein aggregation, further leading to neuronal damage and death^[55][56]^. In addition, histopathological examination of brain tissues from FTD patients shows neuronal loss, microvacuolation, and gliosis; the basal ganglia are often involved, and some cases show neuronal swelling, neuronal loss, and glial cell activation. These findings confirm that the above-enriched nervous system-related biological pathways may be associated with the occurrence and development of FTD^[57]^.

**Figure 1.**
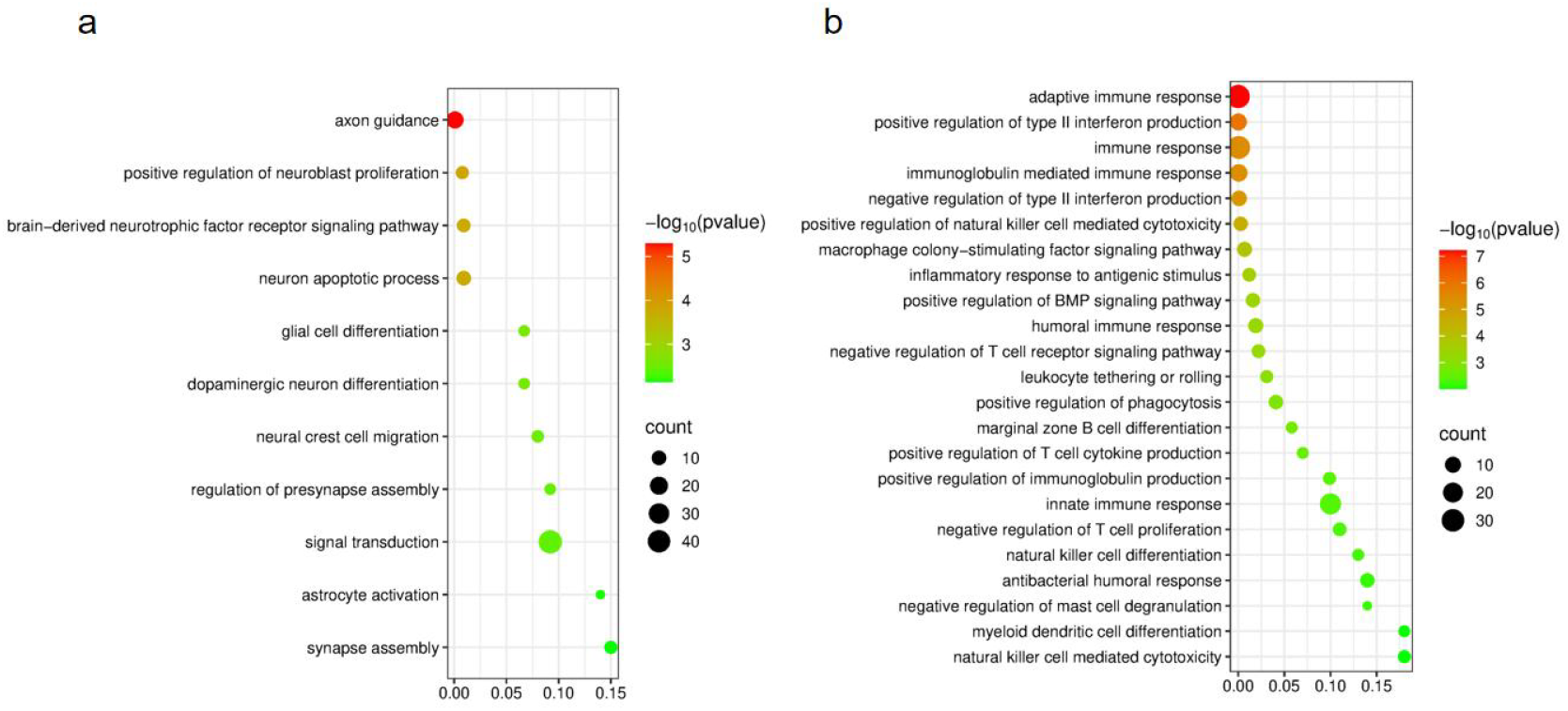
Biological processes enriched by differential proteins between the group with FTD family history without known mutations and the healthy group. a. Pathways related to neurons and nervous system; b. Pathways related to inflammation and immunity.

Among these biological processes, many are related to inflammatory and immune processes, as shown in Figure 1b. Neuronal dysfunction/death, abnormal protein accumulation, and activation of the central immune system play important roles in the pathological progression of FTD. Abnormal protein conformations and accumulation can activate the immune system, thereby inducing neuroinflammation^[58]^. Abnormalities related to neuroinflammation have been found in clinical observations, histological studies, and detection of biomarkers in body fluids such as cerebrospinal fluid from FTD patients^[59]^.

### 3.3 Cluster Analysis of Samples from Individuals with FTD Family History Without Gene Mutations, Carriers with FTD Family History and Gene Mutations, and FTD Patients with MAPT Gene Mutations

Hierarchical cluster analysis was performed on the total proteins of samples from individuals with FTD family history without gene mutations (No. 1, 2, 3, 4, 6), carriers with FTD family history and gene mutations (No. 5), and FTD patients with MAPT gene mutations (No. 7). Proteins with the same or similar expression patterns were clustered, and a total protein cluster analysis map was obtained (Figure 2). As shown in the figure, all 5 samples from individuals with FTD family history without gene mutations clustered together, and healthy individuals except H7 also clustered together. This further confirms that the total proteins of individuals with FTD family history without gene mutations have different characteristics from those of healthy individuals.

**Figure 2.**
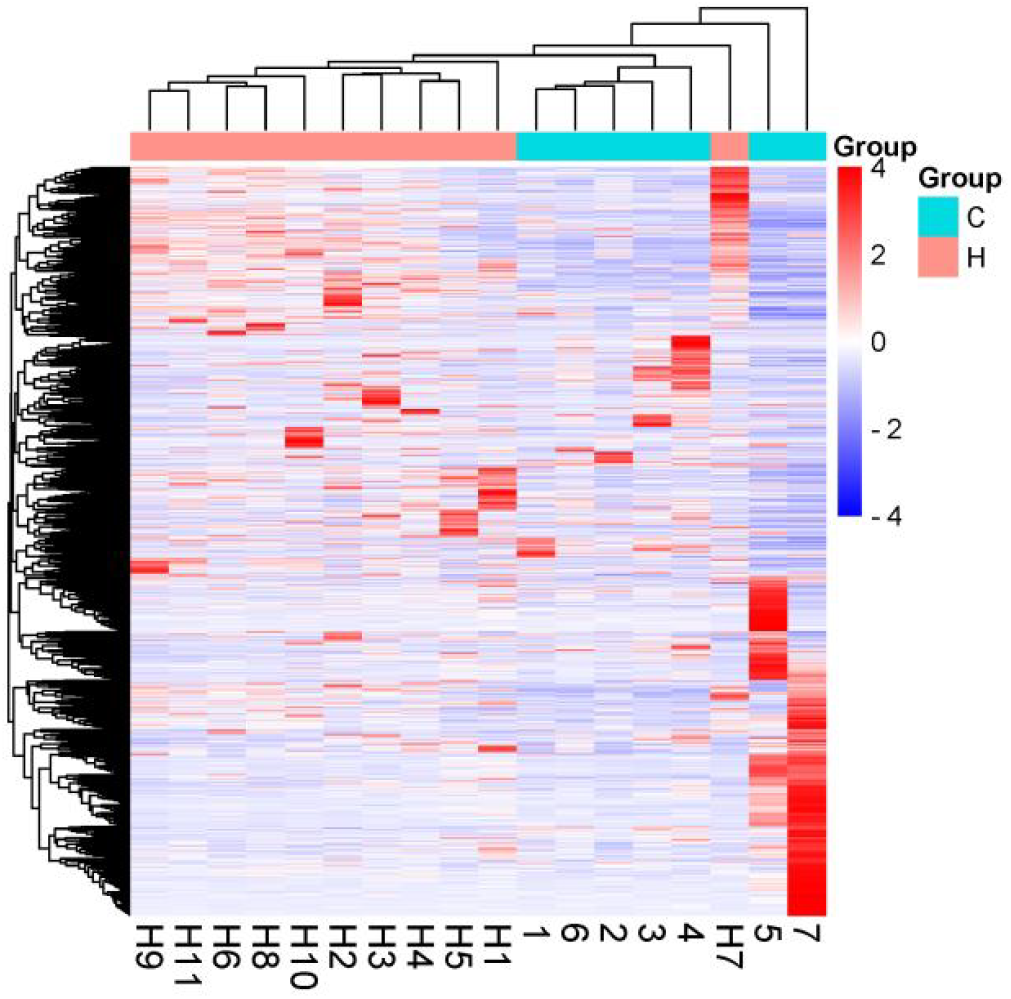
Cluster analysis of samples from individuals with FTD family history without gene mutations (No. 1, 2, 3, 4, 6), carriers with FTD family history and gene mutations (No. 5), and FTD patients with MAPT gene mutations (No. 7).

## 4 Discussion

In this study, 428 differential proteins (P < 0.01, FC ≥ 2.0 or ≤ 0.5) were identified in individuals with a family history of FTD without known mutations through urinary proteomic analysis, among which the significant downregulation of PGRN is particularly critical. As the product encoded by the GRN gene, PGRN haploinsufficiency is one of the core pathogenic mechanisms of FTD^[8]^. Previous studies have shown that GRN mutations can lead to decreased plasma PGRN levels, while this study observed PGRN downregulation in individuals with a family history but without definite mutations, suggesting that even without carrying known mutations, individuals with a family history of FTD may still have PGRN-related pathological cascades^[47]^.

Meanwhile, among the differential proteins, there are various proteins related to neurodegenerative diseases, neurons, or the nervous system, which have undergone significant changes in individuals with a family history of FTD without known mutations. For example, proteins involved in neuronal function or survival such as Integrin beta-3 and Frizzled-5 even showed changes “from presence to absence”. These molecular changes are consistent with the pathological features of neuronal loss and gliosis in the brain tissues of FTD patients, confirming the difference in urinary proteomes between individuals with a family history of FTD without known mutations and healthy individuals, and also demonstrating the sensitivity of the urinary proteome.

Notably, the “maintenance of blood-brain barrier” pathway was enriched among the biological processes (BP) of differential proteins, which may explain the presence of many proteins originally functioning in the brain in urine.

In addition, in the cluster analysis plot (Figure 2), aggregation of members from the same family was observed (No. 1, 2, 3, 4, 6 are members of one family; No. 5, 7 are members of another family). No. 1, 2, 3, 4, 6 are individuals without MAPT gene mutations, while No. 5, 7 carry MAPT mutations. Such genotypic differences may be the core factor driving the respective aggregation of total proteins in the samples. Gene mutations may regulate protein expression patterns, leading to convergent total protein expression characteristics in family members with and without mutations, respectively. Meanwhile, these two families are from different regions (Liaoning Province and Hebei Province). Previous studies have shown that the urinary proteomes of people from different regions in China have significant geographical differences, and region has a great impact on human urinary proteins, which may also affect the aggregation of total proteins in family members in the plot^[60]^_𰀬_

This study has some limitations, including a small sample size and the lack of longitudinal follow-up. Although random grouping verification showed that the randomness of differential proteins was only 4.24%, sample limitations may lead to a certain degree of false positive risk, and the correlation between protein changes and disease progression cannot be clarified. Future studies can expand the sample size and include multicenter cohorts. In addition, longitudinal follow-up can be attempted, combined with imaging techniques, to clarify the association between protein changes and clinical symptoms.

## 5 Conclusion

There are significant differences in the urinary proteomes between individuals with a family history of FTD without known mutations and healthy individuals. This suggests that urinary proteomics has potential in exploring unknown pathogenic factors of FTD, providing a new perspective for deepening disease understanding and optimizing preventive and therapeutic strategies.

## Declarations

## Abbreviations

FTD: Frontotemporal Dementia
PGRN: Progranulin
MAPT: Microtubule - Associated Protein T
GRN: Granulin Gene
FC: Fold change
BP: Biological process

## Ethics approval and consent to participate

The study was conducted in accordance with the Declaration of Helsinki, and approved by the Ethics Committee of Xuanwu Hospital (Clinical Research Review ^[2023]^036).

## Consent for publication

Written informed consent for the publication of the research findings has been obtained from all participants.

## Availability of data and materials

The mass spectrometry proteomics data of this experiment are available in the the project ID of the iProX dataset: IPX0010409000. (https://www.iprox.cn/page/SSV024.html;url=17332128206842Gg6).

## Competing Interests

The authors declare no conflicts of interest.

## Funding

National Key R&D Program of China (2023YFA1801900), Beijing Natural Science Foundation (L246002), Beijing Normal University (11100704).

## Authors’ contributions

YS: responsible for Data Collection and Arrangement, Result Verification and Confirmation, and Paper Writing.

MC & LW: responsible for Providing Experimental Materials and Resources and Data Management.

HW & YS: jointly responsible for Conducting Experimental Operations.

YG: responsible for Paper Revision, Project Management and Coordination, Fundraising and Support, and Data Management.

YG & LW: jointly responsible for Experimental Design.

HW, YS & YG: jointly responsible for Data Analysis.

All authors have reviewed the manuscript.

## Acknowledgements

We appreciate the research of SUN J, WEI J and others. Their available data have helped us draw conclusions. Thanks to Ziyun Shen for her contributions to sample collection.

## Notes

### Competing Interest Statement

The authors have declared no competing interest.

### Summary of Updates

① Adjustments to the overall structure of the article: The original "Results and Discussion" section has been split into two independent sections: "Results" and "Discussion". The "Discussion" section has been established as a separate section, which, on the basis of integrating the results, conducts in-depth analysis of the biological significance in combination with existing literature, and supplements the limitations of the study and future directions. ② Re-analysis of data: The description of samples has been revised (from "unaffected individuals with a family history of frontotemporal dementia" to "unaffected individuals with a family history of frontotemporal dementia without known pathogenic mutations"). A sample that did not belong to the scope of the experimental group (unaffected individuals with a family history of frontotemporal dementia without known pathogenic mutations) was excluded, and the data were re-analyzed. In addition, the description in the abstract has been revised based on the new analysis results.

## References

[1] McKhann GM, Albert MS, Grossman M, et al. Clinical and pathological diagnosis of frontotemporal dementia: report of the Work Group on Frontotemporal Dementia and Pick’s Disease. Arch Neurol 2001;58:1803–9.

[2] Moore KM, Nicholas J, Grossman M, et al. Age at symptom onset and death and disease duration in genetic frontotemporal dementia: an international retrospective cohort study. Lancet Neurol 2020;19 (2):145–156.

[3] Young JJ, Lavakumar M, Tampi D, Balachandran S, Tampi RR. Frontotemporal dementia: latest evidence and clinical implications. Ther Adv Psychopharmacol. 2018 Jan;8(1):33–48. doi: 10.1177/2045125317739818. Epub 2017 Nov 10. PMID: 29344342; PMCID: PMC5761910.

[4] Hodges JR, Piguet O. Progress and Challenges in Frontotemporal Dementia Research: A 20-Year Review. J Alzheimers Dis. 2018;62(3):1467–1480. doi: 10.3233/JAD-171087. PMID: 29504536; PMCID: PMC5870022.

[5] Medina-Rioja R, Gonzalez-Calderon G, Masellis M. Frontotemporal dementia. CMAJ. 2023 Dec 10;195(48):E1660. doi: 10.1503/cmaj.230407. PMID: 38081627; PMCID: PMC10718268.

[6] Greaves CV, Rohrer JD. An update on genetic frontotemporal dementia. J Neurol. 2019 Aug;266(8):2075–2086. doi: 10.1007/s00415-019-09363-4. Epub 2019 May 22. PMID: 31119452; PMCID: PMC6647117.

[7] Seelaar H, Rohrer JD, Pijnenburg YA, Fox NC, van Swieten JC. Clinical, genetic and pathological heterogeneity of frontotemporal dementia: a review. J Neurol Neurosurg Psychiatry. 2011 May;82(5):476–86. doi: 10.1136/jnnp.2010.212225. Epub 2010 Oct 22. PMID: 20971753.

[8] Sevigny J, Uspenskaya O, Heckman LD, Wong LC, Hatch DA, Tewari A, Vandenberghe R, Irwin DJ, Saracino D, Le Ber I, Ahmed R, Rohrer JD, Boxer AL, Boland S, Sheehan P, Brandes A, Burstein SR, Shykind BM, Kamalakaran S, Daniels CW, David Litwack E, Mahoney E, Velaga J, McNamara I, Sondergaard P, Sajjad SA, Kobayashi YM, Abeliovich A, Hefti F. Progranulin AAV gene therapy for frontotemporal dementia: translational studies and phase 1/2 trial interim results. Nat Med. 2024 May;30(5):1406–1415. doi: 10.1038/s41591-024-02973-0. Epub 2024 May 14. PMID: 38745011; PMCID: PMC11108785.

[9] Jicha GA. Medical management of frontotemporal dementias: the importance of the caregiver in symptom assessment and guidance of treatment strategies. J Mol Neurosci. 2011 Nov;45(3):713–23. doi: 10.1007/s12031-011-9558-7. Epub 2011 Jun 7. PMID: 21647712; PMCID: PMC3208136.

[10] Boxer AL, Boeve BF. Frontotemporal dementia treatment: current symptomatic therapies and implications of recent genetic, biochemical, and neuroimaging studies. Alzheimer Dis Assoc Disord. 2007;21:S79–S87.

[11] Tsai RM, Boxer AL. Therapy and clinical trials in frontotemporal dementia: past, present, and future. J Neurochem. 2016 Aug;138 Suppl 1(Suppl 1):211–21. doi: 10.1111/jnc.13640. Epub 2016 Jun 15. PMID: 27306957; PMCID: PMC5217534.

[12] Ntymenou S, Tsantzali I, Kalamatianos T, Voumvourakis KI, Kapaki E, Tsivgoulis G, Stranjalis G, Paraskevas GP. Blood Biomarkers in Frontotemporal Dementia: Review and Meta-Analysis. Brain Sci. 2021 Feb 15;11(2):244. doi: 10.3390/brainsci11020244. PMID: 33672008; PMCID: PMC7919273.

[13] Schoonenboom NS, Reesink FE, Verwey NA, Kester MI, Teunissen CE, van de Ven PM, Pijnenburg YA, Blankenstein MA, Rozemuller AJ, Scheltens P, van der Flier WM. Cerebrospinal fluid markers for differential dementia diagnosis in a large memory clinic cohort. Neurology. 2012 Jan 3;78(1):47–54. doi: 10.1212/WNL.0b013e31823ed0f0. Epub 2011 Dec 14. PMID: 22170879.

[14] Gao Y. Urine-an untapped goldmine for biomarker discovery? Sci China Life Sci. 2013 Dec;56(12):1145–6. doi: 10.1007/s11427-013-4574-1. Epub 2013 Nov 21. PMID: 24271956.

[15] Qin W, Du Z, Gao Y. Collection and preservation of urinary proteins, using a fluff pulp diaper. Sci China Life Sci. 2018 Jun;61(6):671–674. doi: 10.1007/s11427-016-9060-2. Epub 2018 Jan 4. PMID: 29318498.

[16] Jing J, Gao Y. Urine biomarkers in the early stages of diseases: current status and perspective. Discov Med. 2018 Feb;25(136):57-65. PMID: 29579412.

[17] Gao Y. On Research and Translation of Urinary Biomarkers. Adv Exp Med Biol. 2021;1306:101–108. doi: 10.1007/978-3-030-63908-2_7. PMID: 33959908.

[18] Li M, Zhao M, Gao Y. Changes of proteins induced by anticoagulants can be more sensitively detected in urine than in plasma[J]. Sci China Life Sci 2014; 57: 649–56.

[19] Zhang F, Ni Y, Yuan Y, et al. Early urinary candidate biomarker discovery in a rat thioacetamide-induced liver fibrosis model[J]. Sci China Life Sci 2018; 61: 1369–81.

[20] Ni Y, Zhang F, An M, et al. [Changes of urinary proteins in a bacterial meningitis rat model] [J]. Sheng Wu Gong Cheng Xue Bao Chin J Biotechnol 2017; 33: 1145–57.

[21] Wu J, Li X, Zhao M, et al. Early Detection of Urinary Proteome Biomarkers for Effective Early Treatment of Pulmonary Fibrosis in a Rat Model[J]. PROTEOMICS - Clin Appl 2017; 11: 1700103.

[22] Wu J, Guo Z, Gao Y. Dynamic changes of urine proteome in a Walker 256 tumor-bearing rat model[J]. Cancer Med 2017; 6: 2713–22.

[23] Zhang F, Wei J, Li X, et al. Early Candidate Urine Biomarkers for Detecting Alzheimer’s Disease Before Amyloid-β Plaque Deposition in an APP (swe)/PSEN1dE9 Transgenic Mouse Model[J]. J Alzheimers Dis 2018; 66: 613–37.

[24] Rubio-Sánchez R, Ríos-Reina R, Ubeda C. Identification of volatile biomarkers of Trichomonas vaginalis infection in vaginal discharge and urine. Appl Microbiol Biotechnol. 2023 Mar 31. doi: 10.1007/s00253-023-12484-6. Epub ahead of print. PMID: 37000228.

[25] Zhou H, Yuen PS, Pisitkun T,et al. Collection,storage,preservation,and normalization of human urinary exosomesfor biomarker discovery. Kidney Int, 2006,69(8):1471–6.

[26] Sun J, Wei J, Yu H, Sun H, Liu X, Zhang Y, Shao C, Sun W, Zhang J, Gao Y. Urine proteomic characterization of active and recovered COVID-19 patients. bioRxiv; 2023. doi: 10.1101/2023.03.12.532269.PPR:PPR629203.

[27] Lagos-Cabré R, Alvarez A, Kong M, Burgos-Bravo F, Cárdenas A, Rojas-Mancilla E, Pérez-Nuñez R, Herrera-Molina R, Rojas F, Schneider P, Herrera-Marschitz M, Quest AFG, van Zundert B, Leyton L. αβ Integrin regulates astrocyte reactivity. J Neuroinflammation. 2017 Sep 29;14(1):194. doi: 10.1186/s12974-017-0968-5. PMID: 28962574; PMCID: PMC5622429.V3

[28] Liu C, Wang Y, Smallwood PM, Nathans J. An essential role for Frizzled5 in neuronal survival in the parafascicular nucleus of the thalamus. J Neurosci. 2008 May 28;28(22):5641–53. doi: 10.1523/JNEUROSCI.1056-08.2008. PMID: 18509025; PMCID: PMC6670808.

[29] Koszinowski S, Buss K, Kaehlcke K, Krieglstein K. Signaling via the transcriptionally regulated activin receptor 2B is a novel mediator of neuronal cell death during chicken ciliary ganglion development. Int J Dev Neurosci. 2015 Apr;41:98–104. doi: 10.1016/j.ijdevneu.2015.01.006. Epub 2015 Feb 3. PMID: 25660516.

[30] Cheng FY, Huang X, Sarangi A, Ketova T, Cooper MK, Litingtung Y, Chiang C. Widespread contribution of Gdf7 lineage to cerebellar cell types and implications for hedgehog-driven medulloblastoma formation. PLoS One. 2012;7(4):e35541. doi: 10.1371/journal.pone.0035541. Epub 2012 Apr 23. PMID: 22539980; PMCID: PMC3335071.

[31] László ZI, Lele Z. Flying under the radar: CDH2 (N-cadherin), an important hub molecule in neurodevelopmental and neurodegenerative diseases. Front Neurosci. 2022 Sep 23;16:972059. doi: 10.3389/fnins.2022.972059. PMID: 36213737; PMCID: PMC9539934.

[32] Bhumika S, Basalingappa KM, Gopenath TS, Basavaraju S. Glycine encephalopathy. Egypt J Neurol Psychiatr Neurosurg. 2022;58(1):132. doi: 10.1186/s41983-022-00567-6. Epub 2022 Nov 17. PMID: 36415754; PMCID: PMC9672649.

[33] Jin P, Qi D, Cui Y, Lenahan C, Deng S, Tao X. Activation of LRP6 with HLY78 Attenuates Oxidative Stress and Neuronal Apoptosis via GSK3 β/Sirt1/PGC-1 α Pathway after ICH. Oxid Med Cell Longev. 2022 Apr 4;2022:7542468. doi: 10.1155/2022/7542468. PMID: 35419167; PMCID: PMC9001077.

[34] Feng Y, Wang S, Yang D, Zheng W, Xia H, Zhu Q, Wang Z, Hu B, Jiang X, Qin X, Ni C, Pan W, Zhao Y, Pan S, Zhang Y, Song W. Inhibition of IFITM3 in cerebrovascular endothelium alleviates Alzheimer’s-related phenotypes. Alzheimers Dement. 2025 Feb;21(2):e14543. doi: 10.1002/alz.14543. Epub 2025 Jan 14. PMID: 39807629; PMCID: PMC11851164.

[35] Okun E, Mattson MP, Arumugam TV. Involvement of Fc receptors in disorders of the central nervous system. Neuromolecular Med. 2010 Jun;12(2):164–78. doi: 10.1007/s12017-009-8099-5. Epub 2009 Oct 21. PMID: 19844812; PMCID: PMC2878892.

[36] Chauvin S, Sobel A. Neuronal stathmins: a family of phosphoproteins cooperating for neuronal development, plasticity and regeneration. Prog Neurobiol. 2015 Mar;126:1–18. doi: 10.1016/j.pneurobio.2014.09.002. Epub 2014 Oct 16. PMID: 25449700.

[37] Adhihetty PJ, Beal MF. Creatine and its potential therapeutic value for targeting cellular energy impairment in neurodegenerative diseases. Neuromolecular Med. 2008;10(4):275–90. doi: 10.1007/s12017-008-8053-y. Epub 2008 Nov 13. PMID: 19005780; PMCID: PMC2886719.

[38] Gómez-Nicola D, Valle-Argos B, Pallas-Bazarra N, Nieto-Sampedro M. Interleukin-15 regulates proliferation and self-renewal of adult neural stem cells. Mol Biol Cell. 2011 Jun 15;22(12):1960–70. doi: 10.1091/mbc.E11-01-0053. Epub 2011 Apr 20. PMID: 21508317; PMCID: PMC3113763.

[39] Kim J, Kim S, Kim H, Hwang IW, Bae S, Karki S, Kim D, Ogelman R, Bang G, Kim JY, Kajander T, Um JW, Oh WC, Ko J. MDGA1 negatively regulates amyloid precursor protein-mediated synapse inhibition in the hippocampus. Proc Natl Acad Sci U S A. 2022 Jan 25;119(4):e2115326119. doi: 10.1073/pnas.2115326119. Erratum in: Proc Natl Acad Sci U S A. 2024 Jun 25;121(26):e2410758121. doi: 10.1073/pnas.2410758121. PMID: 35074912; PMCID: PMC8795569.

[40] Sharma S, Behl T, Kumar A, Sehgal A, Singh S, Sharma N, Bhatia S, Al-Harrasi A, Bungau S. Targeting Endothelin in Alzheimer’s Disease: A Promising Therapeutic Approach. Biomed Res Int. 2021 Sep 6;2021:7396580. doi: 10.1155/2021/7396580. PMID: 34532504; PMCID: PMC8440097.

[41] Bloch-Gallego E, Anderson DI. Key role of Rho GTPases in motor disorders associated with neurodevelopmental pathologies. Mol Psychiatry. 2023 Jan;28(1):118–126. doi: 10.1038/s41380-022-01702-8. Epub 2022 Aug 2. Erratum in: Mol Psychiatry. 2023 Jan;28(1):515. doi: 10.1038/s41380-022-01847-6. PMID: 35918397.

[42] Zhao Y, Wang Y, Yang J, Wang X, Zhao Y, Zhang X, Zhang YW. Sorting nexin 12 interacts with BACE1 and regulates BACE1-mediated APP processing. Mol Neurodegener. 2012 Jun 18;7:30. doi: 10.1186/1750-1326-7-30. PMID: 22709416; PMCID: PMC3439308.

[43] Ceballos-Picot I, Mockel L, Potier MC, Dauphinot L, Shirley TL, Torero-Ibad R, Fuchs J, Jinnah HA. Hypoxanthine-guanine phosphoribosyl transferase regulates early developmental programming of dopamine neurons: implications for Lesch-Nyhan disease pathogenesis. Hum Mol Genet. 2009 Jul 1;18(13):2317–27. doi: 10.1093/hmg/ddp164. Epub 2009 Apr 2. PMID: 19342420; PMCID: PMC2694685.

[44] Gaweda-Walerych K, Aragona V, Lodato S, Sitek EJ, Narożańska E, Buratti E. Progranulin deficiency in the brain: the interplay between neuronal and non-neuronal cells. Transl Neurodegener. 2025 Apr 16;14(1):18. doi: 10.1186/s40035-025-00475-8. PMID: 40234992; PMCID: PMC12001433.

[45] Huang M, Modeste E, Dammer E, Merino P, Taylor G, Duong DM, Deng Q, Holler CJ, Gearing M, Dickson D, Seyfried NT, Kukar T. Network analysis of the progranulin-deficient mouse brain proteome reveals pathogenic mechanisms shared in human frontotemporal dementia caused by GRN mutations. Acta Neuropathol Commun. 2020 Oct 7;8(1):163. doi: 10.1186/s40478-020-01037-x. PMID: 33028409; PMCID: PMC7541308.

[46] Mukherjee O, Wang J, Gitcho M, Chakraverty S, Taylor-Reinwald L, Shears S, Kauwe JS, Norton J, Levitch D, Bigio EH, Hatanpaa KJ, White CL, Morris JC, Cairns NJ, Goate A. Molecular characterization of novel progranulin (GRN) mutations in frontotemporal dementia. Hum Mutat. 2008 Apr;29(4):512–21. doi: 10.1002/humu.20681. PMID: 18183624; PMCID: PMC2756561.

[47] Sellami L, Rucheton B, Ben Younes I, Camuzat A, Saracino D, Rinaldi D, Epelbaum S, Azuar C, Levy R, Auriacombe S, Hannequin D, Pariente J, Barbier M, Boutoleau-Bretonnière C, Couratier P, Pasquier F, Deramecourt V, Sauvée M, Sarazin M, Lagarde J, Roué-Jagot C, Forlani S, Jornea L, David I; French Research Network on FTLD/FTLD-ALS; PREVDEMALS and Predict-PGRN Groups; LeGuern E, Dubois B, Brice A, Clot F, Lamari F, Le Ber I. Plasma progranulin levels for frontotemporal dementia in clinical practice: a 10-year French experience. Neurobiol Aging. 2020 Jul;91:167.e1-167.e9. doi: 10.1016/j.neurobiolaging.2020.02.014. Epub 2020 Feb 21. PMID: 32171590.

[48] Mahoney ER, Dumitrescu L, Moore AM, Cambronero FE, De Jager PL, Koran MEI, Petyuk VA, Robinson RAS, Goyal S, Schneider JA, Bennett DA, Jefferson AL, Hohman TJ. Brain expression of the vascular endothelial growth factor gene family in cognitive aging and alzheimer’s disease. Mol Psychiatry. 2021 Mar;26(3):888–896. doi: 10.1038/s41380-019-0458-5. Epub 2019 Jul 22. PMID: 31332262; PMCID: PMC6980445.

[49] Lorenzl S, Buerger K, Hampel H, Beal MF. Profiles of matrix metalloproteinases and their inhibitors in plasma of patients with dementia. Int Psychogeriatr. 2008 Feb;20(1):67–76. doi: 10.1017/S1041610207005790. Epub 2007 Aug 15. PMID: 17697439.

[50] Tuna G, Yener GG, Oktay G, İşlekel GH, Kİrkalİ FG. Evaluation of Matrix Metalloproteinase-2 (MMP-2) and -9 (MMP-9) and Their Tissue Inhibitors (TIMP-1 and TIMP-2) in Plasma from Patients with Neurodegenerative Dementia. J Alzheimers Dis. 2018;66(3):1265–1273. doi: 10.3233/JAD-180752. PMID: 30412498.

[51] Wennström M, Nielsen HM. Cell adhesion molecules in Alzheimer’s disease. Degener Neurol Neuromuscul Dis. 2012 Jul 4;2:65–77. doi: 10.2147/DNND.S19829. PMID: 30890880; PMCID: PMC6065556.

[52] Akoudad S, Wolters FJ, Viswanathan A, de Bruijn RF, van der Lugt A, Hofman A, Koudstaal PJ, Ikram MA, Vernooij MW. Association of Cerebral Microbleeds With Cognitive Decline and Dementia. JAMA Neurol. 2016 Aug 1;73(8):934–43. doi: 10.1001/jamaneurol.2016.1017. PMID: 27271785; PMCID: PMC5966721.

[53] Ghasemi M, Keyhanian K, Douthwright C. Glial Cell Dysfunction in C9orf72-Related Amyotrophic Lateral Sclerosis and Frontotemporal Dementia. Cells. 2021 Jan 28;10(2):249. doi: 10.3390/cells10020249. PMID: 33525344; PMCID: PMC7912327.

[54] Marinelli S, Marrone MC, Di Domenico M, Marinelli S. Endocannabinoid signaling in microglia. Glia. 2023 Jan;71(1):71–90. doi: 10.1002/glia.24281. Epub 2022 Oct 12. PMID: 36222019.

[55] Valori CF, Sulmona C, Brambilla L, Rossi D. Astrocytes: Dissecting Their Diverse Roles in Amyotrophic Lateral Sclerosis and Frontotemporal Dementia. Cells. 2023 May 23;12(11):1450. doi: 10.3390/cells12111450. PMID: 37296571; PMCID: PMC10252425.

[56] McCauley ME, Baloh RH. Inflammation in ALS/FTD pathogenesis. Acta Neuropathol. 2019 May;137(5):715–730. doi: 10.1007/s00401-018-1933-9. Epub 2018 Nov 21. PMID: 30465257; PMCID: PMC6482122.

[57] Cagnin A, Rossor M, Sampson EL, Mackinnon T, Banati RB. In vivo detection of microglial activation in frontotemporal dementia. Ann Neurol. 2004 Dec;56(6):894–7. doi: 10.1002/ana.20332. PMID: 15562429.

[58] Borrego-Écija S, Pérez-Millan A, Antonell A, Fort-Aznar L, Kaya-Tilki E, León-Halcón A, Lladó A, Molina-Porcel L, Balasa M, Juncà-Parella J, Vitorica J, Venero JL, Deierborg T, Boza-Serrano A, Sánchez-Valle R. Galectin-3 is upregulated in frontotemporal dementia patients with subtype specificity. Alzheimers Dement. 2024 Mar;20(3):1515–1526. doi: 10.1002/alz.13536. Epub 2023 Nov 29. PMID: 38018380; PMCID: PMC10984429.

[59] Bright F, Werry EL, Dobson-Stone C, Piguet O, Ittner LM, Halliday GM, Hodges JR, Kiernan MC, Loy CT, Kassiou M, Kril JJ. Neuroinflammation in frontotemporal dementia. Nat Rev Neurol. 2019 Sep;15(9):540–555. doi: 10.1038/s41582-019-0231-z. Epub 2019 Jul 19. PMID: 31324897.

[60] Wu JQ, Qin WW, Pan L, Wang XR, Zhang B, Shan GL, Gao YH. Regional Differences of the Urinary Proteomes in Healthy Chinese Individuals. Chin Med Sci J. 2019 Sep 30;34(3):157–167. doi: 10.24920/003504. PMID: 31601298.

