## Appendix Table for "The urinary proteome of individuals with a family history of frontotemporal dementia (FTD) but without known mutations differs from that of healthy controls"

Table 1 Differential protein information screened by FC≥2.0 or ≤0.5 and P<0.01 between the group with a family history of frontotemporal dementia and no known mutations and the healthy group

| Uniprot ID | Protein names | Fold change | P value | Trend |
| --- | --- | --- | --- | --- |
| Q13467 | Frizzled-5 | 0 | 2.57E-05 | ↓ |
| P05106 | Integrin beta-3 | 0 | 2.08E-04 | ↓ |
| Q6UXH9 | Inactive serine protease PAMR1 | 0 | 4.82E-04 | ↓ |
| Q9NRR1 | Cytokine-like protein 1 | 0 | 1.17E-03 | ↓ |
| Q9NQ34 | Transmembrane protein 9B | 0 | 1.47E-03 | ↓ |
| Q13705 | Activin receptor type-2B | 0 | 2.08E-03 | ↓ |
| Q7Z4P5 | Growth/differentiation factor 7 | 0 | 3.11E-03 | ↓ |
| P30486 | HLA class I histocompatibility antigen, B alpha chain | 0 | 5.57E-03 | ↓ |
| P26012 | Integrin beta-8 | 0.00114775 | 3.22E-05 | ↓ |
| A0A0C4DH35 | Probable non-functional immunoglobulin heavy variable 3-35 | 0.003278932 | 3.90E-06 | ↓ |
| P01782 | Immunoglobulin heavy variable 3-9 | 0.004192089 | 9.33E-05 | ↓ |
| A0A087WW87 | Immunoglobulin kappa variable 2-40 | 0.004206211 | 1.75E-03 | ↓ |
| Q86UP0 | Cadherin-24 | 0.004366636 | 3.19E-03 | ↓ |
| P23434 | Glycine cleavage system H protein, mitochondrial | 0.005513261 | 3.48E-04 | ↓ |
| P0DP01 | Immunoglobulin heavy variable 1-8 | 0.014288546 | 3.84E-05 | ↓ |
| O75581 | Low-density lipoprotein receptor-related protein 6 | 0.017474301 | 4.04E-03 | ↓ |
| Q01628 | Interferon-induced transmembrane protein 3 | 0.01844234 | 9.17E-05 | ↓ |
| Q7KYR7 | Butyrophilin subfamily 2 member A1 | 0.018457592 | 4.14E-04 | ↓ |
| P35556 | Fibrillin-2 [Cleaved into: Placensin] | 0.0217965 | 6.30E-03 | ↓ |
| O75015 | Low affinity immunoglobulin gamma Fc region receptor III-B | 0.026270901 | 1.44E-03 | ↓ |
| Q96MM6 | Heat shock 70 kDa protein 12B | 0.026908408 | 3.78E-03 | ↓ |
| P06310 | Immunoglobulin kappa variable 2-30 | 0.030212044 | 7.60E-03 | ↓ |
| P24394 | Interleukin-4 receptor subunit alpha | 0.031487263 | 5.17E-03 | ↓ |
| Q9NQS3 | Nectin-3 | 0.0322231 | 2.55E-05 | ↓ |
| Q6B0K9 | Hemoglobin subunit mu | 0.033304355 | 1.17E-05 | ↓ |
| P23083 | Immunoglobulin heavy variable 1-2 | 0.033580957 | 8.01E-04 | ↓ |
| P35968 | Vascular endothelial growth factor receptor 2 | 0.037206217 | 4.96E-05 | ↓ |
| O60635 | Tetraspanin-1 | 0.037323421 | 9.33E-06 | ↓ |
| O75493 | Carbonic anhydrase-related protein 11 | 0.038563178 | 9.32E-03 | ↓ |
| O43653 | Prostate stem cell antigen | 0.039833282 | 9.05E-03 | ↓ |
| P07998 | Ribonuclease pancreatic | 0.040986707 | 6.58E-07 | ↓ |
| Q86Y78 | Ly6/PLAUR domain-containing protein 6 | 0.041324487 | 4.82E-05 | ↓ |
| P62081 | Small ribosomal subunit protein eS7 | 0.04464649 | 5.67E-03 | ↓ |
| Q9BZG1 | Ras-related protein Rab-34 | 0.046596535 | 1.26E-03 | ↓ |
| Q6IAA8 | Ragulator complex protein LAMTOR1 | 0.0466007 | 2.43E-05 | ↓ |
| Q96A25 | Transmembrane protein 106A | 0.048219488 | 7.79E-04 | ↓ |
| P05451 | Lithostathine-1-alpha | 0.052584545 | 5.17E-03 | ↓ |
| P0DI83 | Ras-related protein Rab-34, isoform NARR | 0.058503967 | 5.37E-03 | ↓ |
| P40199 | Cell adhesion molecule CEACAM6 | 0.059444074 | 4.27E-05 | ↓ |
| P55084 | Trifunctional enzyme subunit beta, mitochondrial | 0.059466482 | 2.44E-03 | ↓ |
| Q96C90 | Protein phosphatase 1 regulatory subunit 14B | 0.060593478 | 4.09E-03 | ↓ |
| P35625 | Metalloproteinase inhibitor 3 | 0.062687539 | 4.32E-03 | ↓ |
| Q8ND76 | Cyclin-Y | 0.063467037 | 6.69E-05 | ↓ |
| P08174 | Complement decay-accelerating factor | 0.065058741 | 2.36E-03 | ↓ |
| Q8NBJ9 | SID1 transmembrane family member 2 | 0.065972095 | 4.07E-03 | ↓ |
| P48048 | ATP-sensitive inward rectifier potassium channel 1 | 0.066927174 | 1.27E-03 | ↓ |
| P30531 | Sodium- and chloride-dependent GABA transporter 1 | 0.069341357 | 7.18E-04 | ↓ |
| P78325 | Disintegrin and metalloproteinase domain-containing protein 8 | 0.070543218 | 8.45E-03 | ↓ |
| P40939 | Trifunctional enzyme subunit alpha, mitochondrial | 0.071070683 | 4.32E-03 | ↓ |
| Q8N357 | Solute carrier family 35 member F6 | 0.07123198 | 9.82E-03 | ↓ |
| Q99732 | Lipopolysaccharide-induced tumor necrosis factor-alpha factor | 0.071802862 | 8.04E-04 | ↓ |
| P13224 | Platelet glycoprotein Ib beta chain | 0.073294944 | 3.47E-04 | ↓ |
| Q9NRY6 | Phospholipid scramblase 3 | 0.076088948 | 8.58E-03 | ↓ |
| Q9BY32 | Inosine triphosphate pyrophosphatase | 0.077628002 | 5.23E-05 | ↓ |
| Q9UGN4 | CMRF35-like molecule 8 | 0.080488103 | 1.76E-05 | ↓ |
| P23327 | Sarcoplasmic reticulum histidine-rich calcium-binding protein | 0.081675566 | 2.21E-03 | ↓ |
| P47895 | Retinaldehyde dehydrogenase 3 | 0.083278719 | 6.39E-03 | ↓ |
| P55884 | Eukaryotic translation initiation factor 3 subunit B | 0.083797147 | 4.54E-05 | ↓ |
| P16109 | P-selectin | 0.085407414 | 1.28E-03 | ↓ |
| Q5TFQ8 | Signal-regulatory protein beta-1 isoform 3 | 0.086347381 | 1.15E-03 | ↓ |
| Q9P0T7 | Proton-transporting V-type ATPase complex assembly regulator TMEM9 | 0.089203796 | 3.06E-06 | ↓ |
| Q96FX2 | Diphthamide biosynthesis protein 3 | 0.090548605 | 7.28E-03 | ↓ |
| O43866 | CD5 antigen-like | 0.094046628 | 3.28E-04 | ↓ |
| P07307 | Asialoglycoprotein receptor 2 | 0.096796341 | 8.22E-06 | ↓ |
| P41439 | Folate receptor gamma | 0.098583207 | 3.08E-03 | ↓ |
| Q03403 | Trefoil factor 2 | 0.099741686 | 6.32E-04 | ↓ |
| O15067 | Phosphoribosylformylglycinamidine synthase | 0.09976529 | 2.38E-04 | ↓ |
| P02749 | Beta-2-glycoprotein 1 | 0.103090525 | 1.03E-04 | ↓ |
| P42658 | A-type potassium channel modulatory protein DPP6 | 0.107606166 | 2.56E-03 | ↓ |
| A0A075B6R2 | Immunoglobulin heavy variable 4-4 | 0.109220867 | 8.84E-03 | ↓ |
| P20594 | Atrial natriuretic peptide receptor 2 | 0.109708088 | 8.17E-03 | ↓ |
| O75094 | Slit homolog 3 protein | 0.110489995 | 2.58E-05 | ↓ |
| Q5VT99 | Leucine-rich repeat-containing protein 38 | 0.117823743 | 2.30E-03 | ↓ |
| Q9BXR6 | Complement factor H-related protein 5 | 0.1189915 | 6.84E-03 | ↓ |
| Q96GP6 | Scavenger receptor class F member 2 | 0.120922538 | 7.01E-04 | ↓ |
| Q9Y2I2 | Netrin-G1 | 0.12263749 | 1.30E-04 | ↓ |
| Q00577 | Transcriptional activator protein Pur-alpha | 0.123056632 | 5.17E-03 | ↓ |
| P0DJD8 | Pepsin A-3 | 0.123081378 | 5.19E-04 | ↓ |
| Q8N1G4 | Leucine-rich repeat-containing protein 47 | 0.126146847 | 8.04E-04 | ↓ |
| Q16853 | Amine oxidase [copper-containing] 3 | 0.128896975 | 8.83E-04 | ↓ |
| P25311 | Zinc-alpha-2-glycoprotein | 0.131993525 | 5.20E-03 | ↓ |
| P14174 | Macrophage migration inhibitory factor | 0.132713703 | 2.80E-04 | ↓ |
| Q96J42 | Thioredoxin domain-containing protein 15 | 0.133387264 | 1.92E-03 | ↓ |
| P35555 | Fibrillin-1 [Cleaved into: Asprosin] | 0.135949111 | 4.64E-06 | ↓ |
| P02458 | Collagen alpha-1 | 0.140088485 | 6.58E-03 | ↓ |
| Q8N114 | Protein shisa-5 | 0.140137825 | 1.97E-03 | ↓ |
| P04114 | Apolipoprotein B-100 | 0.142050963 | 7.66E-04 | ↓ |
| Q6UXB4 | C-type lectin domain family 4 member G | 0.145927642 | 1.52E-07 | ↓ |
| P01130 | Low-density lipoprotein receptor | 0.146336737 | 2.12E-04 | ↓ |
| O75629 | Protein CREG1 | 0.147630704 | 1.74E-05 | ↓ |
| P35659 | Protein DEK | 0.151383052 | 5.28E-03 | ↓ |
| Q86VB7 | Scavenger receptor cysteine-rich type 1 protein M130 | 0.153650345 | 4.49E-05 | ↓ |
| Q8NFT8 | Delta and Notch-like epidermal growth factor-related receptor | 0.154685522 | 1.08E-04 | ↓ |
| O43921 | Ephrin-A2 | 0.154865974 | 2.00E-04 | ↓ |
| P06276 | Cholinesterase | 0.156642462 | 3.78E-04 | ↓ |
| Q9H7M9 | V-type immunoglobulin domain-containing suppressor of T-cell activation | 0.156945084 | 2.16E-03 | ↓ |
| Q9BXS4 | Transmembrane protein 59 | 0.15759673 | 4.17E-03 | ↓ |
| P05090 | Apolipoprotein D | 0.157783831 | 1.03E-06 | ↓ |
| Q8TEM1 | Nuclear pore membrane glycoprotein 210 | 0.15833812 | 7.83E-03 | ↓ |
| A1L4H1 | Soluble scavenger receptor cysteine-rich domain-containing protein SSC5D | 0.158507231 | 2.82E-05 | ↓ |
| Q14508 | WAP four-disulfide core domain protein 2 | 0.158566537 | 8.13E-05 | ↓ |
| Q14162 | Scavenger receptor class F member 1 | 0.159325856 | 2.06E-05 | ↓ |
| P01834 | Immunoglobulin kappa constant | 0.161761203 | 1.28E-03 | ↓ |
| P01589 | Interleukin-2 receptor subunit alpha | 0.162476998 | 6.76E-06 | ↓ |
| Q04721 | Neurogenic locus notch homolog protein 2 | 0.164288079 | 7.65E-07 | ↓ |
| O60241 | Adhesion G protein-coupled receptor B2 | 0.165142909 | 1.83E-04 | ↓ |
| Q66K79 | Carboxypeptidase Z | 0.165890434 | 1.64E-04 | ↓ |
| Q9NVG8 | TBC1 domain family member 13 | 0.166355302 | 7.87E-03 | ↓ |
| P41222 | Prostaglandin-H2 D-isomerase | 0.166911943 | 2.67E-03 | ↓ |
| P16562 | Cysteine-rich secretory protein 2 | 0.167105386 | 5.15E-05 | ↓ |
| P06731 | Cell adhesion molecule CEACAM5 | 0.167750091 | 7.50E-03 | ↓ |
| O00187 | Mannan-binding lectin serine protease 2 | 0.168837013 | 1.03E-03 | ↓ |
| P48960 | Adhesion G protein-coupled receptor E5 | 0.170499197 | 2.41E-03 | ↓ |
| P02452 | Collagen alpha-1 | 0.170661303 | 3.08E-03 | ↓ |
| P02753 | Retinol-binding protein 4 | 0.175979656 | 1.11E-03 | ↓ |
| Q86VP1 | Tax1-binding protein 1 | 0.176915692 | 4.04E-03 | ↓ |
| P31785 | Cytokine receptor common subunit gamma | 0.177845242 | 8.71E-04 | ↓ |
| Q14050 | Collagen alpha-3 | 0.178420059 | 6.33E-04 | ↓ |
| O75023 | Leukocyte immunoglobulin-like receptor subfamily B member 5 | 0.182628473 | 2.40E-03 | ↓ |
| Q8NHL6 | Leukocyte immunoglobulin-like receptor subfamily B member 1 | 0.184261011 | 7.39E-04 | ↓ |
| P49184 | Deoxyribonuclease-1-like 1 | 0.18459112 | 7.27E-03 | ↓ |
| Q86TY3 | Armadillo-like helical domain-containing protein 4 | 0.187935251 | 9.96E-03 | ↓ |
| Q96H15 | T-cell immunoglobulin and mucin domain-containing protein 4 | 0.188706612 | 3.60E-04 | ↓ |
| Q9NPF0 | CD320 antigen | 0.191402834 | 1.59E-06 | ↓ |
| A6NDG6 | Glycerol-3-phosphate phosphatase | 0.194291939 | 1.40E-06 | ↓ |
| P47972 | Neuronal pentraxin-2 | 0.195343253 | 3.94E-05 | ↓ |
| Q13201 | Multimerin-1 | 0.196363576 | 7.79E-04 | ↓ |
| P13688 | Cell adhesion molecule CEACAM1 | 0.19707471 | 1.03E-06 | ↓ |
| P57087 | Junctional adhesion molecule B | 0.197127687 | 6.60E-03 | ↓ |
| Q9P232 | Contactin-3 | 0.197263431 | 8.29E-03 | ↓ |
| O94919 | Endonuclease domain-containing 1 protein | 0.19847285 | 1.45E-03 | ↓ |
| Q9H3S7 | Tyrosine-protein phosphatase non-receptor type 23 | 0.198932911 | 1.58E-03 | ↓ |
| Q6ZUM4 | Rho GTPase-activating protein 27 | 0.199786702 | 2.17E-04 | ↓ |
| Q8N126 | Cell adhesion molecule 3 | 0.201067377 | 2.97E-03 | ↓ |
| A0A0B4J1V6 | Immunoglobulin heavy variable 3-73 | 0.205432579 | 8.29E-07 | ↓ |
| Q99983 | Osteomodulin | 0.206586981 | 6.28E-03 | ↓ |
| Q16378 | Proline-rich protein 4 | 0.206686566 | 1.87E-03 | ↓ |
| P24752 | Acetyl-CoA acetyltransferase, mitochondrial | 0.209223973 | 3.33E-03 | ↓ |
| P12830 | Cadherin-1 | 0.209571855 | 1.88E-03 | ↓ |
| P61769 | Beta-2-microglobulin [Cleaved into: Beta-2-microglobulin form pI 5.3] | 0.210198944 | 8.63E-03 | ↓ |
| Q8IZ21 | Phosphatase and actin regulator 4 | 0.21195471 | 3.80E-03 | ↓ |
| Q99466 | Neurogenic locus notch homolog protein 4 | 0.212258076 | 6.67E-03 | ↓ |
| Q07507 | Dermatopontin | 0.212264854 | 1.52E-04 | ↓ |
| P13598 | Intercellular adhesion molecule 2 | 0.212700791 | 3.71E-06 | ↓ |
| Q86SJ2 | Amphoterin-induced protein 2 | 0.212890824 | 5.94E-04 | ↓ |
| Q6UWL2 | Sushi domain-containing protein 1 | 0.216370651 | 2.12E-03 | ↓ |
| Q10588 | ADP-ribosyl cyclase/cyclic ADP-ribose hydrolase 2 | 0.219245944 | 2.28E-04 | ↓ |
| Q8N2U0 | Transmembrane protein 256 | 0.221654777 | 7.19E-03 | ↓ |
| O00182 | Galectin-9 | 0.221896608 | 1.42E-03 | ↓ |
| Q9NP58 | ATP-binding cassette sub-family B member 6 | 0.224716384 | 1.63E-03 | ↓ |
| Q5VWZ2 | Lysophospholipase-like protein 1 | 0.226805768 | 3.38E-03 | ↓ |
| O95971 | CD160 antigen | 0.227854025 | 2.21E-05 | ↓ |
| P09972 | Fructose-bisphosphate aldolase C | 0.230236137 | 1.88E-04 | ↓ |
| P36980 | Complement factor H-related protein 2 | 0.230417477 | 1.16E-03 | ↓ |
| Q6UY11 | Protein delta homolog 2 | 0.230514268 | 2.94E-05 | ↓ |
| A0A0C4DH25 | Immunoglobulin kappa variable 3D-20 | 0.232263126 | 2.89E-03 | ↓ |
| P01876 | Immunoglobulin heavy constant alpha 1 | 0.23270023 | 5.33E-05 | ↓ |
| Q5SRE7 | Phytanoyl-CoA dioxygenase domain-containing protein 1 | 0.234122168 | 2.42E-03 | ↓ |
| P25189 | Myelin protein P0 | 0.234632807 | 1.36E-03 | ↓ |
| A0PJK1 | Sodium/mannose cotransporter SLC5A10 | 0.237724786 | 1.05E-03 | ↓ |
| Q14353 | Guanidinoacetate N-methyltransferase | 0.238872445 | 4.62E-04 | ↓ |
| Q9HCU0 | Endosialin | 0.239087869 | 1.34E-05 | ↓ |
| Q08431 | Lactadherin | 0.240982877 | 3.43E-03 | ↓ |
| P01764 | Immunoglobulin heavy variable 3-23 | 0.241153429 | 1.41E-03 | ↓ |
| O15431 | High affinity copper uptake protein 1 | 0.242675012 | 2.63E-05 | ↓ |
| P62269 | Small ribosomal subunit protein uS13 | 0.242836531 | 4.51E-03 | ↓ |
| Q96DR8 | Mucin-like protein 1 | 0.242885344 | 3.19E-03 | ↓ |
| Q07812 | Apoptosis regulator BAX | 0.246825937 | 2.77E-03 | ↓ |
| Q13291 | Signaling lymphocytic activation molecule | 0.247305467 | 6.86E-05 | ↓ |
| P05452 | Tetranectin | 0.25155342 | 3.47E-05 | ↓ |
| A0A0C4DH32 | Immunoglobulin heavy variable 3-20 | 0.251941852 | 2.49E-03 | ↓ |
| P08887 | Interleukin-6 receptor subunit alpha | 0.252956854 | 4.87E-05 | ↓ |
| Q15762 | CD226 antigen | 0.254164735 | 7.91E-06 | ↓ |
| P98172 | Ephrin-B1 | 0.254951743 | 3.53E-04 | ↓ |
| P13473 | Lysosome-associated membrane glycoprotein 2 | 0.255004151 | 1.17E-04 | ↓ |
| Q9NRX4 | 14 kDa phosphohistidine phosphatase | 0.255376263 | 3.72E-03 | ↓ |
| Q6UXI9 | Nephronectin | 0.257090985 | 4.26E-04 | ↓ |
| Q9UNN8 | Endothelial protein C receptor | 0.26158643 | 2.98E-04 | ↓ |
| O43854 | EGF-like repeat and discoidin I-like domain-containing protein 3 | 0.261855048 | 4.08E-03 | ↓ |
| Q96PX8 | SLIT and NTRK-like protein 1 | 0.262134265 | 5.67E-04 | ↓ |
| P02768 | Albumin | 0.264511021 | 2.35E-06 | ↓ |
| O75594 | Peptidoglycan recognition protein 1 | 0.266826208 | 2.92E-05 | ↓ |
| P32942 | Intercellular adhesion molecule 3 | 0.26745504 | 6.43E-07 | ↓ |
| Q9Y5X3 | Sorting nexin-5 | 0.268305766 | 5.48E-03 | ↓ |
| Q9UM47 | Neurogenic locus notch homolog protein 3 | 0.268508691 | 2.11E-03 | ↓ |
| P61970 | Nuclear transport factor 2 | 0.268580116 | 7.85E-05 | ↓ |
| Q9HCH3 | Copine-5 | 0.268975591 | 1.96E-03 | ↓ |
| Q9Y3B3 | Transmembrane emp24 domain-containing protein 7 | 0.269034281 | 8.83E-04 | ↓ |
| P19652 | Alpha-1-acid glycoprotein 2 | 0.269126561 | 1.15E-03 | ↓ |
| Q9Y6N7 | Roundabout homolog 1 | 0.269398597 | 6.00E-04 | ↓ |
| P15692 | Vascular endothelial growth factor A, long form | 0.269609426 | 5.12E-03 | ↓ |
| P06870 | Kallikrein-1 | 0.270899349 | 1.93E-05 | ↓ |
| P08519 | Apolipoprotein | 0.271681553 | 7.64E-04 | ↓ |
| P04216 | Thy-1 membrane glycoprotein | 0.271848094 | 9.21E-04 | ↓ |
| P28799 | Progranulin | 0.273410965 | 7.70E-04 | ↓ |
| Q9HBH0 | Rho-related GTP-binding protein RhoF | 0.274571409 | 7.79E-03 | ↓ |
| Q8N2S1 | Latent-transforming growth factor beta-binding protein 4 | 0.274650799 | 6.54E-03 | ↓ |
| Q9NZN1 | Interleukin-1 receptor accessory protein-like 1 | 0.275177588 | 2.96E-03 | ↓ |
| Q04756 | Hepatocyte growth factor activator | 0.276543846 | 1.61E-05 | ↓ |
| O00337 | Sodium/nucleoside cotransporter 1 | 0.276553984 | 8.75E-04 | ↓ |
| Q9UKY0 | Prion-like protein doppel | 0.276992112 | 7.02E-04 | ↓ |
| Q9H461 | Frizzled-8 | 0.27770636 | 6.75E-05 | ↓ |
| P02763 | Alpha-1-acid glycoprotein 1 | 0.277741144 | 6.57E-03 | ↓ |
| Q8NFY4 | Semaphorin-6D | 0.282031162 | 3.49E-03 | ↓ |
| Q14031 | Collagen alpha-6 | 0.284391778 | 5.93E-03 | ↓ |
| P26992 | Ciliary neurotrophic factor receptor subunit alpha | 0.28446729 | 8.80E-04 | ↓ |
| Q8IZP9 | Adhesion G-protein coupled receptor G2 | 0.285104157 | 3.06E-03 | ↓ |
| Q16363 | Laminin subunit alpha-4 | 0.290230577 | 5.61E-04 | ↓ |
| P02787 | Serotransferrin | 0.290945598 | 5.76E-05 | ↓ |
| P01210 | Proenkephalin-A [Cleaved into: Synenkephalin; Met-enkephalin | 0.291230872 | 7.28E-03 | ↓ |
| P08571 | Monocyte differentiation antigen CD14 | 0.292707126 | 3.23E-03 | ↓ |
| P53801 | Pituitary tumor-transforming gene 1 protein-interacting protein | 0.292892738 | 3.50E-04 | ↓ |
| P28908 | Tumor necrosis factor receptor superfamily member 8 | 0.293995846 | 6.47E-04 | ↓ |
| Q9NQ36 | Signal peptide, CUB and EGF-like domain-containing protein 2 | 0.295457077 | 5.64E-04 | ↓ |
| P07359 | Platelet glycoprotein Ib alpha chain | 0.297352075 | 3.69E-03 | ↓ |
| Q99835 | Protein smoothened | 0.298960143 | 2.84E-04 | ↓ |
| P41181 | Aquaporin-2 | 0.300144761 | 1.50E-03 | ↓ |
| Q86TD4 | Sarcalumenin | 0.300722187 | 4.42E-03 | ↓ |
| Q12907 | Vesicular integral-membrane protein VIP36 | 0.300931742 | 7.85E-04 | ↓ |
| O95865 | Putative hydrolase DDAH2 | 0.302668256 | 1.03E-04 | ↓ |
| P46531 | Neurogenic locus notch homolog protein 1 | 0.303470368 | 7.98E-05 | ↓ |
| Q96HD1 | Protein disulfide isomerase CRELD1 | 0.307019294 | 2.03E-03 | ↓ |
| Q9H6X2 | Anthrax toxin receptor 1 | 0.308532186 | 6.56E-03 | ↓ |
| O43895 | Xaa-Pro aminopeptidase 2 | 0.309210857 | 2.56E-05 | ↓ |
| P02462 | Collagen alpha-1 | 0.310324125 | 1.27E-03 | ↓ |
| P08709 | Coagulation factor VII | 0.3120223 | 8.77E-03 | ↓ |
| Q9Y5Y7 | Lymphatic vessel endothelial hyaluronic acid receptor 1 | 0.312213226 | 5.48E-03 | ↓ |
| O95398 | Rap guanine nucleotide exchange factor 3 | 0.313810443 | 1.33E-03 | ↓ |
| P00734 | Prothrombin | 0.314722221 | 5.16E-06 | ↓ |
| Q8N3J6 | Cell adhesion molecule 2 | 0.314995094 | 6.23E-05 | ↓ |
| O95998 | Interleukin-18-binding protein | 0.315379707 | 1.33E-03 | ↓ |
| P0DOX2 | Immunoglobulin alpha-2 heavy chain | 0.317776087 | 3.09E-05 | ↓ |
| Q9UP38 | Frizzled-1 | 0.318909719 | 1.92E-03 | ↓ |
| Q9HAR2 | Adhesion G protein-coupled receptor L3 | 0.319308107 | 5.79E-04 | ↓ |
| Q8TDQ0 | Hepatitis A virus cellular receptor 2 | 0.320386705 | 2.43E-06 | ↓ |
| Q9H8L6 | Multimerin-2 | 0.321269214 | 7.11E-07 | ↓ |
| P02790 | Hemopexin | 0.321313274 | 1.37E-07 | ↓ |
| O95490 | Adhesion G protein-coupled receptor L2 | 0.321345647 | 1.09E-06 | ↓ |
| Q96GW7 | Brevican core protein | 0.323083603 | 1.96E-03 | ↓ |
| Q12805 | EGF-containing fibulin-like extracellular matrix protein 1 | 0.323294299 | 9.04E-04 | ↓ |
| P21802 | Fibroblast growth factor receptor 2 | 0.323911409 | 6.98E-05 | ↓ |
| P31997 | Cell adhesion molecule CEACAM8 | 0.324319809 | 4.18E-03 | ↓ |
| P08637 | Low affinity immunoglobulin gamma Fc region receptor III-A | 0.324869993 | 7.33E-04 | ↓ |
| P98095 | Fibulin-2 | 0.328043784 | 1.36E-04 | ↓ |
| P10912 | Growth hormone receptor | 0.329094658 | 1.29E-04 | ↓ |
| P11597 | Cholesteryl ester transfer protein | 0.33126665 | 1.07E-03 | ↓ |
| P16422 | Epithelial cell adhesion molecule | 0.332126383 | 3.01E-06 | ↓ |
| Q16719 | Kynureninase | 0.333235991 | 5.61E-03 | ↓ |
| Q9UBG0 | C-type mannose receptor 2 | 0.334477138 | 9.32E-05 | ↓ |
| Q9Y6Q6 | Tumor necrosis factor receptor superfamily member 11A | 0.33559421 | 1.45E-03 | ↓ |
| Q5VY43 | Platelet endothelial aggregation receptor 1 | 0.337906409 | 2.65E-03 | ↓ |
| P04217 | Alpha-1B-glycoprotein | 0.338465535 | 9.42E-03 | ↓ |
| O95336 | 6-phosphogluconolactonase | 0.338550642 | 1.08E-04 | ↓ |
| Q9UBD6 | Ammonium transporter Rh type C | 0.339110999 | 7.68E-04 | ↓ |
| Q8WVV5 | Butyrophilin subfamily 2 member A2 | 0.343213985 | 4.86E-07 | ↓ |
| Q08334 | Interleukin-10 receptor subunit beta | 0.344564105 | 4.02E-04 | ↓ |
| P0DOX5 | Immunoglobulin gamma-1 heavy chain | 0.345349592 | 2.86E-08 | ↓ |
| O43280 | Trehalase | 0.346506318 | 4.74E-03 | ↓ |
| Q96F10 | Thialysine N-epsilon-acetyltransferase | 0.348089028 | 3.95E-05 | ↓ |
| Q6UXG3 | CMRF35-like molecule 9 | 0.348161861 | 6.38E-05 | ↓ |
| P15260 | Interferon gamma receptor 1 | 0.348168441 | 8.68E-03 | ↓ |
| Q9NZP8 | Complement C1r subcomponent-like protein | 0.350970154 | 1.17E-06 | ↓ |
| P13987 | CD59 glycoprotein | 0.35116614 | 9.33E-04 | ↓ |
| Q93070 | Ecto-ADP-ribosyltransferase 4 | 0.351939931 | 9.93E-03 | ↓ |
| Q6EMK4 | Vasorin | 0.353291039 | 7.81E-04 | ↓ |
| P16066 | Atrial natriuretic peptide receptor 1 | 0.353492505 | 8.12E-05 | ↓ |
| P04746 | Pancreatic alpha-amylase | 0.353520815 | 5.20E-04 | ↓ |
| P15328 | Folate receptor alpha | 0.354012354 | 5.87E-04 | ↓ |
| Q6FHJ7 | Secreted frizzled-related protein 4 | 0.356636835 | 2.77E-03 | ↓ |
| Q9P121 | Neurotrimin | 0.357447029 | 1.02E-03 | ↓ |
| Q13873 | Bone morphogenetic protein receptor type-2 | 0.358268646 | 2.89E-04 | ↓ |
| Q9BXJ7 | Protein amnionless [Cleaved into: Soluble protein amnionless] | 0.358323428 | 2.29E-05 | ↓ |
| P09564 | T-cell antigen CD7 | 0.362055601 | 6.47E-03 | ↓ |
| Q7Z3B1 | Neuronal growth regulator 1 | 0.363435813 | 1.43E-03 | ↓ |
| Q5ZPR3 | CD276 antigen | 0.363719418 | 9.82E-04 | ↓ |
| O14498 | Immunoglobulin superfamily containing leucine-rich repeat protein | 0.363751084 | 3.06E-05 | ↓ |
| P40197 | Platelet glycoprotein V | 0.365146379 | 1.13E-04 | ↓ |
| A6NL88 | Protein shisa-7 | 0.365598673 | 5.80E-03 | ↓ |
| Q15375 | Ephrin type-A receptor 7 | 0.366655227 | 8.11E-06 | ↓ |
| O15162 | Phospholipid scramblase 1 | 0.368473622 | 1.02E-03 | ↓ |
| P29972 | Aquaporin-1 | 0.369114782 | 5.86E-03 | ↓ |
| Q96B86 | Repulsive guidance molecule A | 0.369133563 | 1.24E-03 | ↓ |
| Q86T13 | C-type lectin domain family 14 member A | 0.369499362 | 3.51E-05 | ↓ |
| Q8N474 | Secreted frizzled-related protein 1 | 0.371283876 | 4.72E-03 | ↓ |
| Q9H9K5 | Endogenous retroviral envelope protein HEMO | 0.372447024 | 1.56E-04 | ↓ |
| Q9UNE0 | Tumor necrosis factor receptor superfamily member EDAR | 0.37266039 | 3.96E-03 | ↓ |
| P01591 | Immunoglobulin J chain | 0.376048773 | 6.67E-04 | ↓ |
| O94973 | AP-2 complex subunit alpha-2 | 0.376136475 | 8.15E-03 | ↓ |
| Q16827 | Receptor-type tyrosine-protein phosphatase O | 0.377681559 | 3.37E-03 | ↓ |
| P02766 | Transthyretin | 0.379363478 | 3.08E-03 | ↓ |
| P21583 | Kit ligand | 0.3812868 | 5.62E-04 | ↓ |
| Q9UIB8 | SLAM family member 5 | 0.382011624 | 1.92E-04 | ↓ |
| P26718 | NKG2-D type II integral membrane protein | 0.382963879 | 1.22E-03 | ↓ |
| Q01415 | N-acetylgalactosamine kinase | 0.382974548 | 2.92E-03 | ↓ |
| P02765 | Alpha-2-HS-glycoprotein | 0.38299508 | 6.33E-03 | ↓ |
| Q8TCT8 | Signal peptide peptidase-like 2A | 0.383052268 | 6.82E-03 | ↓ |
| P41240 | Tyrosine-protein kinase CSK | 0.383399153 | 9.27E-03 | ↓ |
| P16112 | Aggrecan core protein | 0.384844133 | 4.39E-04 | ↓ |
| Q8NBR0 | Tumor protein p53-inducible protein 13 | 0.385020697 | 3.94E-03 | ↓ |
| Q5UCC4 | ER membrane protein complex subunit 10 | 0.386066925 | 2.32E-03 | ↓ |
| P04156 | Major prion protein | 0.3864521 | 4.26E-03 | ↓ |
| P62979 | Ubiquitin-ribosomal protein eS31 fusion protein | 0.387931013 | 4.38E-04 | ↓ |
| P04745 | Alpha-amylase 1A | 0.388952355 | 2.39E-04 | ↓ |
| Q8WZ75 | Roundabout homolog 4 | 0.389218757 | 5.46E-04 | ↓ |
| P01833 | Polymeric immunoglobulin receptor | 0.391302595 | 2.30E-04 | ↓ |
| P20933 | N | 0.393741541 | 3.25E-03 | ↓ |
| P00740 | Coagulation factor IX | 0.395515873 | 4.36E-06 | ↓ |
| O95967 | EGF-containing fibulin-like extracellular matrix protein 2 | 0.396139846 | 7.98E-04 | ↓ |
| P60033 | CD81 antigen | 0.396393697 | 6.95E-03 | ↓ |
| Q9Y281 | Cofilin-2 | 0.396558123 | 4.53E-03 | ↓ |
| P07204 | Thrombomodulin | 0.397159409 | 1.75E-04 | ↓ |
| Q9UJ99 | Cadherin-22 | 0.39815959 | 2.32E-03 | ↓ |
| P35443 | Thrombospondin-4 | 0.399197354 | 3.86E-04 | ↓ |
| P04406 | Glyceraldehyde-3-phosphate dehydrogenase | 0.399574385 | 7.62E-04 | ↓ |
| P01859 | Immunoglobulin heavy constant gamma 2 | 0.400264959 | 2.46E-04 | ↓ |
| P25325 | 3-mercaptopyruvate sulfurtransferase | 0.40044589 | 1.12E-04 | ↓ |
| Q92563 | Testican-2 | 0.401453403 | 1.65E-03 | ↓ |
| P82980 | Retinol-binding protein 5 | 0.401532752 | 8.60E-04 | ↓ |
| Q96H20 | Vacuolar-sorting protein SNF8 | 0.402077563 | 1.25E-03 | ↓ |
| O75509 | Tumor necrosis factor receptor superfamily member 21 | 0.402498781 | 8.39E-04 | ↓ |
| Q9HCN6 | Platelet glycoprotein VI | 0.402875283 | 1.07E-03 | ↓ |
| O75339 | Cartilage intermediate layer protein 1 | 0.403456378 | 2.37E-03 | ↓ |
| P00505 | Aspartate aminotransferase, mitochondrial | 0.403623636 | 3.62E-03 | ↓ |
| Q53RD9 | Fibulin-7 | 0.403833194 | 3.25E-03 | ↓ |
| Q9NZ53 | Podocalyxin-like protein 2 | 0.404122536 | 1.29E-03 | ↓ |
| P61916 | NPC intracellular cholesterol transporter 2 | 0.406637037 | 1.50E-03 | ↓ |
| Q6UX15 | Layilin | 0.406738975 | 2.51E-03 | ↓ |
| Q92692 | Nectin-2 | 0.40694508 | 3.16E-04 | ↓ |
| Q496F6 | CMRF35-like molecule 2 | 0.407796522 | 1.62E-03 | ↓ |
| Q96KN2 | Beta-Ala-His dipeptidase | 0.409049435 | 1.30E-03 | ↓ |
| Q14894 | Ketimine reductase mu-crystallin | 0.409072129 | 3.05E-03 | ↓ |
| Q6GTX8 | Leukocyte-associated immunoglobulin-like receptor 1 | 0.409427316 | 9.55E-04 | ↓ |
| Q9HCP0 | Casein kinase I isoform gamma-1 | 0.409767392 | 4.74E-04 | ↓ |
| O14594 | Neurocan core protein | 0.410144378 | 3.27E-03 | ↓ |
| Q96RW7 | Hemicentin-1 | 0.410640198 | 1.35E-05 | ↓ |
| P25940 | Collagen alpha-3 | 0.410926577 | 6.27E-04 | ↓ |
| O00592 | Podocalyxin | 0.411628874 | 3.68E-03 | ↓ |
| O15197 | Ephrin type-B receptor 6 | 0.412322568 | 3.30E-05 | ↓ |
| Q8IZA0 | Dyslexia-associated protein KIAA0319-like protein | 0.413055594 | 1.20E-04 | ↓ |
| P11279 | Lysosome-associated membrane glycoprotein 1 | 0.413681075 | 4.22E-03 | ↓ |
| P14151 | L-selectin | 0.413954912 | 2.90E-03 | ↓ |
| P31689 | DnaJ homolog subfamily A member 1 | 0.414916536 | 4.12E-03 | ↓ |
| A0A0B4J1Y9 | Immunoglobulin heavy variable 3-72 | 0.416065408 | 6.26E-03 | ↓ |
| P41217 | OX-2 membrane glycoprotein | 0.417182616 | 9.08E-04 | ↓ |
| P19022 | Cadherin-2 | 0.41731033 | 1.95E-03 | ↓ |
| P12259 | Coagulation factor V | 0.418184512 | 2.33E-04 | ↓ |
| Q8IYS5 | Osteoclast-associated immunoglobulin-like receptor | 0.421574461 | 6.65E-04 | ↓ |
| Q96GD0 | Chronophin | 0.422448781 | 5.82E-03 | ↓ |
| P26927 | Hepatocyte growth factor-like protein | 0.422783868 | 4.70E-04 | ↓ |
| P00748 | Coagulation factor XII | 0.424766958 | 8.26E-03 | ↓ |
| P11362 | Fibroblast growth factor receptor 1 | 0.425854228 | 2.77E-04 | ↓ |
| Q96CS7 | Pleckstrin homology domain-containing family B member 2 | 0.426256013 | 5.03E-03 | ↓ |
| P43121 | Cell surface glycoprotein MUC18 | 0.42871229 | 8.84E-05 | ↓ |
| Q8IUK5 | Plexin domain-containing protein 1 | 0.431534392 | 1.22E-03 | ↓ |
| Q9UKZ9 | Procollagen C-endopeptidase enhancer 2 | 0.432321058 | 4.41E-03 | ↓ |
| Q9BY67 | Cell adhesion molecule 1 | 0.434446794 | 7.15E-04 | ↓ |
| Q9UKU9 | Angiopoietin-related protein 2 | 0.435399506 | 4.63E-03 | ↓ |
| P32241 | Vasoactive intestinal polypeptide receptor 1 | 0.437291561 | 6.54E-04 | ↓ |
| O00161 | Synaptosomal-associated protein 23 | 0.438479584 | 4.43E-04 | ↓ |
| P01042 | Kininogen-1 | 0.439187183 | 2.02E-05 | ↓ |
| Q9BXP8 | Pappalysin-2 | 0.439243075 | 5.26E-03 | ↓ |
| Q9Y287 | Integral membrane protein 2B | 0.439374136 | 1.97E-05 | ↓ |
| P80370 | Protein delta homolog 1 | 0.439679631 | 3.51E-03 | ↓ |
| Q9BRN9 | TM2 domain-containing protein 3 | 0.439910787 | 1.90E-03 | ↓ |
| P17927 | Complement receptor type 1 | 0.440299389 | 7.82E-03 | ↓ |
| Q92626 | Peroxidasin homolog | 0.442840546 | 3.46E-03 | ↓ |
| P00450 | Ceruloplasmin | 0.448633479 | 1.66E-03 | ↓ |
| P30530 | Tyrosine-protein kinase receptor UFO | 0.450682946 | 1.74E-03 | ↓ |
| Q9GZX9 | Twisted gastrulation protein homolog 1 | 0.452721142 | 5.35E-03 | ↓ |
| P42330 | Aldo-keto reductase family 1 member C3 | 0.455852845 | 2.02E-03 | ↓ |
| P05556 | Integrin beta-1 | 0.456244751 | 1.43E-07 | ↓ |
| Q9UK23 | N-acetylglucosamine-1-phosphodiester alpha-N-acetylglucosaminidase | 0.459546592 | 2.35E-03 | ↓ |
| Q8N307 | Mucin-20 | 0.463317596 | 2.57E-03 | ↓ |
| Q96AP7 | Endothelial cell-selective adhesion molecule | 0.46377773 | 6.24E-03 | ↓ |
| P36888 | Receptor-type tyrosine-protein kinase FLT3 | 0.464410133 | 2.73E-03 | ↓ |
| P02649 | Apolipoprotein E | 0.465687613 | 2.55E-03 | ↓ |
| P98160 | Basement membrane-specific heparan sulfate proteoglycan core protein | 0.467086044 | 2.26E-04 | ↓ |
| O95834 | Echinoderm microtubule-associated protein-like 2 | 0.468408664 | 6.18E-03 | ↓ |
| P01133 | Pro-epidermal growth factor | 0.470368701 | 3.49E-03 | ↓ |
| P27797 | Calreticulin | 0.470406752 | 1.25E-03 | ↓ |
| Q9NZZ3 | Charged multivesicular body protein 5 | 0.4710442 | 3.98E-03 | ↓ |
| Q9H9P2 | Chondrolectin | 0.474063038 | 4.31E-03 | ↓ |
| Q10471 | Polypeptide N-acetylgalactosaminyltransferase 2 | 0.474346381 | 3.84E-03 | ↓ |
| Q8WWZ8 | Oncoprotein-induced transcript 3 protein | 0.475445183 | 4.85E-04 | ↓ |
| P21709 | Ephrin type-A receptor 1 | 0.482918031 | 1.38E-03 | ↓ |
| P10643 | Complement component C7 | 0.484656413 | 7.36E-04 | ↓ |
| P54802 | Alpha-N-acetylglucosaminidase | 0.485956282 | 4.52E-04 | ↓ |
| Q9NZV1 | Cysteine-rich motor neuron 1 protein | 0.486447118 | 7.54E-05 | ↓ |
| O94910 | Adhesion G protein-coupled receptor L1 | 0.486964049 | 4.55E-03 | ↓ |
| Q06418 | Tyrosine-protein kinase receptor TYRO3 | 0.489862362 | 8.26E-03 | ↓ |
| P07203 | Glutathione peroxidase 1 | 0.490233889 | 2.83E-03 | ↓ |
| P02774 | Vitamin D-binding protein | 0.490518938 | 3.98E-05 | ↓ |
| Q92890 | Ubiquitin recognition factor in ER-associated degradation protein 1 | 0.491290693 | 9.17E-03 | ↓ |
| Q14767 | Latent-transforming growth factor beta-binding protein 2 | 0.491706383 | 8.10E-04 | ↓ |
| P13591 | Neural cell adhesion molecule 1 | 0.491812533 | 7.90E-03 | ↓ |
| P11047 | Laminin subunit gamma-1 | 0.492406106 | 7.17E-04 | ↓ |
| P54826 | Growth arrest-specific protein 1 | 0.494088671 | 1.60E-04 | ↓ |
| P05067 | Amyloid-beta precursor protein | 0.496983131 | 8.63E-05 | ↓ |
| P31939 | Bifunctional purine biosynthesis protein ATIC | 0.498987327 | 1.14E-03 | ↓ |
| O75368 | Adapter SH3BGRL | 2.008690294 | 8.81E-03 | ↑ |
| P60953 | Cell division control protein 42 homolog | 2.134968422 | 8.60E-03 | ↑ |
| P11413 | Glucose-6-phosphate 1-dehydrogenase | 2.171450922 | 9.38E-04 | ↑ |
| P80303 | Nucleobindin-2 | 2.308256198 | 2.40E-04 | ↑ |
| Q9Y5I2 | Protocadherin alpha-10 | 2.382837211 | 2.04E-03 | ↑ |
| P20339 | Ras-related protein Rab-5A | 2.418272736 | 5.19E-04 | ↑ |
| O75348 | V-type proton ATPase subunit G 1 | 2.442693681 | 9.52E-03 | ↑ |
| Q8NI35 | InaD-like protein | 2.525639738 | 3.93E-03 | ↑ |
| P35475 | Alpha-L-iduronidase | 2.572050695 | 6.89E-03 | ↑ |
| P00492 | Hypoxanthine-guanine phosphoribosyltransferase | 2.636819012 | 8.18E-03 | ↑ |
| Q9UMY4 | Sorting nexin-12 | 2.829118948 | 3.69E-03 | ↑ |
| Q9Y3B8 | Oligoribonuclease, mitochondrial | 2.854295325 | 2.92E-03 | ↑ |
| P84095 | Rho-related GTP-binding protein RhoG | 2.87999873 | 5.48E-03 | ↑ |
| Q9H3Z4 | DnaJ homolog subfamily C member 5 | 2.967484834 | 2.43E-03 | ↑ |
| O43681 | ATPase GET3 | 3.430697148 | 3.45E-03 | ↑ |
| P12081 | Histidine--tRNA ligase, cytoplasmic | 3.44649458 | 1.44E-04 | ↑ |
| Q9H3K6 | BolA-like protein 2 | 3.448032927 | 7.38E-03 | ↑ |
| P14138 | Endothelin-3 | 3.519859507 | 7.04E-03 | ↑ |
| P52943 | Cysteine-rich protein 2 | 3.788360332 | 1.44E-03 | ↑ |
| Q8NFP4 | MAM domain-containing glycosylphosphatidylinositol anchor protein 1 | 4.127538855 | 5.30E-04 | ↑ |
| Q13261 | Interleukin-15 receptor subunit alpha | 4.39874634 | 2.93E-03 | ↑ |
| P54577 | Tyrosine--tRNA ligase, cytoplasmic | 4.728219431 | 2.11E-03 | ↑ |
| Q8N5J2 | Ubiquitin carboxyl-terminal hydrolase MINDY-1 | 5.954260168 | 1.74E-06 | ↑ |
| P12532 | Creatine kinase U-type, mitochondrial | 6.769300018 | 9.66E-05 | ↑ |
| O00534 | von Willebrand factor A domain-containing protein 5A | 7.252552388 | 1.96E-04 | ↑ |
| Q2TAA2 | Isoamyl acetate-hydrolyzing esterase 1 homolog | 10.28191409 | 8.46E-03 | ↑ |
| P35247 | Pulmonary surfactant-associated protein D | 15.09223155 | 1.51E-03 | ↑ |
| P16949 | Stathmin | 18.25924665 | 2.32E-03 | ↑ |
| P62857 | Small ribosomal subunit protein eS28 | 21.28955763 | 2.95E-03 | ↑ |

Table 2 BP (Biological Process) information enriched from differential proteins between the group with a family history of frontotemporal dementia and no known mutations and the healthy group

| Term | P-Value |
| --- | --- |
| cell adhesion | 1.20E-18 |
| heterophilic cell-cell adhesion via plasma membrane cell adhesion molecules | 5.20E-10 |
| angiogenesis | 6.90E-09 |
| blood coagulation | 8.10E-09 |
| homophilic cell adhesion via plasma membrane adhesion molecules | 8.60E-09 |
| cell surface receptor signaling pathway | 1.70E-08 |
| collagen-activated tyrosine kinase receptor signaling pathway | 4.90E-08 |
| adaptive immune response | 5.90E-08 |
| symbiont entry into host cell | 1.00E-07 |
| zymogen activation | 6.20E-07 |
| positive regulation of type II interferon production | 1.10E-06 |
| glomerular capillary formation | 3.40E-06 |
| hepatocyte growth factor receptor signaling pathway | 3.50E-06 |
| immune response | 3.70E-06 |
| immunoglobulin mediated immune response | 4.00E-06 |
| positive regulation of cell population proliferation | 4.10E-06 |
| axon guidance | 5.00E-06 |
| negative regulation of type II interferon production | 6.70E-06 |
| cellular response to tumor cell | 7.80E-06 |
| cell-cell adhesion | 8.00E-06 |
| positive regulation of smooth muscle cell differentiation | 8.80E-06 |
| atrial septum morphogenesis | 1.30E-05 |
| blood coagulation, intrinsic pathway | 1.50E-05 |
| positive regulation of ERK1 and ERK2 cascade | 1.60E-05 |
| epidermal growth factor receptor signaling pathway | 1.80E-05 |
| ephrin receptor signaling pathway | 1.80E-05 |
| atrioventricular node development | 2.70E-05 |
| positive regulation of natural killer cell mediated cytotoxicity | 2.80E-05 |
| left/right axis specification | 3.50E-05 |
| positive regulation of tumor necrosis factor production | 3.90E-05 |
| receptor-mediated endocytosis | 4.00E-05 |
| intracellular copper ion homeostasis | 4.70E-05 |
| vascular endothelial growth factor signaling pathway | 6.40E-05 |
| negative regulation of gene expression | 6.70E-05 |
| complement activation, classical pathway | 7.20E-05 |
| insulin-like growth factor receptor signaling pathway | 7.50E-05 |
| positive regulation of Ras protein signal transduction | 7.90E-05 |
| pulmonary valve morphogenesis | 7.90E-05 |
| vascular endothelial growth factor receptor-1 signaling pathway | 9.50E-05 |
| podocyte development | 1.00E-04 |
| intrahepatic bile duct development | 1.00E-04 |
| regulation of osteoclast development | 1.00E-04 |
| cholangiocyte proliferation | 1.00E-04 |
| platelet-derived growth factor receptor-alpha signaling pathway | 1.10E-04 |
| BMP signaling pathway | 1.20E-04 |
| cell migration | 1.20E-04 |
| Kit signaling pathway | 1.20E-04 |
| macrophage colony-stimulating factor signaling pathway | 1.20E-04 |
| positive regulation of platelet activation | 1.40E-04 |
| positive regulation of neuroblast proliferation | 1.50E-04 |
| brain-derived neurotrophic factor receptor signaling pathway | 1.80E-04 |
| platelet-derived growth factor receptor-beta signaling pathway | 1.80E-04 |
| neuron apoptotic process | 1.80E-04 |
| intracellular iron ion homeostasis | 1.90E-04 |
| positive regulation of phosphatidylinositol 3-kinase/protein kinase B signal transduction | 1.90E-04 |
| negative regulation of blood coagulation | 2.00E-04 |
| vascular endothelial growth factor receptor-2 signaling pathway | 2.00E-04 |
| acute-phase response | 2.30E-04 |
| cell migration involved in sprouting angiogenesis | 2.30E-04 |
| placenta blood vessel development | 2.70E-04 |
| inflammatory response to antigenic stimulus | 2.80E-04 |
| ciliary body morphogenesis | 3.50E-04 |
| proximal tubule development | 3.50E-04 |
| positive regulation of BMP signaling pathway | 3.90E-04 |
| humoral immune response | 4.70E-04 |
| negative regulation of T cell receptor signaling pathway | 5.50E-04 |
| bone remodeling | 5.70E-04 |
| positive regulation of gene expression | 6.10E-04 |
| platelet activation | 6.30E-04 |
| hemopoiesis | 6.30E-04 |
| adherens junction organization | 6.40E-04 |
| vascular wound healing | 8.20E-04 |
| bone trabecula formation | 8.20E-04 |
| leukocyte tethering or rolling | 8.60E-04 |
| positive regulation of bone resorption | 1.00E-03 |
| cell fate determination | 1.00E-03 |
| phagocytosis | 1.10E-03 |
| extracellular matrix organization | 1.10E-03 |
| vagina development | 1.10E-03 |
| embryonic eye morphogenesis | 1.10E-03 |
| regulation of protein metabolic process | 1.10E-03 |
| hepatocyte proliferation | 1.10E-03 |
| Notch signaling pathway | 1.20E-03 |
| animal organ regeneration | 1.30E-03 |
| positive regulation of phagocytosis | 1.30E-03 |
| negative regulation of cell-cell adhesion mediated by cadherin | 1.60E-03 |
| lysosomal lumen acidification | 1.80E-03 |
| embryonic limb morphogenesis | 1.80E-03 |
| positive regulation of osteoblast differentiation | 2.00E-03 |
| positive regulation of cell migration | 2.00E-03 |
| positive regulation of MAPK cascade | 2.00E-03 |
| regulation of peptidyl-tyrosine phosphorylation | 2.00E-03 |
| marginal zone B cell differentiation | 2.00E-03 |
| artery morphogenesis | 2.10E-03 |
| glial cell differentiation | 2.40E-03 |
| dopaminergic neuron differentiation | 2.40E-03 |
| morphogenesis of an epithelial sheet | 2.60E-03 |
| positive regulation of vascular endothelial growth factor signaling pathway | 2.60E-03 |
| positive regulation of T cell cytokine production | 2.60E-03 |
| non-canonical Wnt signaling pathway | 2.80E-03 |
| peptidyl-tyrosine phosphorylation | 3.10E-03 |
| neural crest cell migration | 3.10E-03 |
| lymphangiogenesis | 3.30E-03 |
| regulation of presynapse assembly | 3.60E-03 |
| positive regulation of endothelial cell migration | 3.60E-03 |
| signal transduction | 3.70E-03 |
| calcium-dependent cell-cell adhesion via plasma membrane cell adhesion molecules | 3.70E-03 |
| response to lipopolysaccharide | 4.00E-03 |
| substrate adhesion-dependent cell spreading | 4.10E-03 |
| positive regulation of immunoglobulin production | 4.10E-03 |
| positive regulation of endothelial cell proliferation | 4.20E-03 |
| innate immune response | 4.40E-03 |
| endocytosis | 4.40E-03 |
| positive regulation of epithelial cell proliferation | 4.50E-03 |
| positive regulation of osteoclast differentiation | 4.60E-03 |
| positive regulation of epithelial tube formation | 4.70E-03 |
| negative regulation of T cell proliferation | 4.80E-03 |
| positive regulation of blood coagulation | 4.90E-03 |
| complement activation, alternative pathway | 4.90E-03 |
| animal organ morphogenesis | 5.10E-03 |
| heart looping | 5.20E-03 |
| natural killer cell differentiation | 5.80E-03 |
| maintenance of blood-brain barrier | 6.50E-03 |
| antibacterial humoral response | 6.70E-03 |
| astrocyte activation | 6.90E-03 |
| positive regulation of membrane protein ectodomain proteolysis | 6.90E-03 |
| positive regulation of viral life cycle | 6.90E-03 |
| negative regulation of mast cell degranulation | 6.90E-03 |
| positive regulation of cellular extravasation | 6.90E-03 |
| synapse assembly | 7.20E-03 |
| positive regulation of peptidyl-tyrosine phosphorylation | 7.20E-03 |
| fibroblast growth factor receptor signaling pathway | 7.50E-03 |
| apoptotic cell clearance | 8.00E-03 |
| negative regulation of anoikis | 8.00E-03 |
| negative regulation of tumor necrosis factor production | 8.60E-03 |
| ureteric bud development | 8.80E-03 |
| skeletal system development | 8.80E-03 |
| myeloid dendritic cell differentiation | 9.30E-03 |
| positive regulation of keratinocyte proliferation | 9.30E-03 |
| glomerular filtration | 9.30E-03 |
| regulation of complement activation | 9.50E-03 |
| lymphocyte proliferation | 9.50E-03 |
| lipoprotein catabolic process | 9.50E-03 |
| positive regulation of smoothened signaling pathway | 9.60E-03 |
| natural killer cell mediated cytotoxicity | 9.60E-03 |
| positive regulation of angiogenesis | 9.80E-03 |
